# Global transmission architecture of VIM carbapenemases reveals host-specific dissemination strategies

**DOI:** 10.64898/2026.08.16.745130

**Authors:** Miaoshan Luo, Ni Li, Jingjie Song, Qingqing Zhi, Ruibin Lai, Minling Wang, Minhong Wang, Ge Wang, Mingxiao Chen, Cong Shen, Qiang Zhou

## Abstract

Carbapenemase-producing Gram-negative bacteria carrying Verona integron-encoded metallo-β-lactamase (VIM) pose a persistent global threat, yet the mechanisms driving their worldwide dissemination remain poorly resolved. We analysed 5,617 *bla*_VIM_-positive genomes collected from 73 countries or regions across six continents between 1999 and 2025 to reconstruct the global epidemiology and transmission architecture of VIM carbapenemases. Forty VIM variants were identified across 16 bacterial genera, revealing marked host preferences and temporal shifts. VIM-2 dominated global circulation, whereas VIM-1 and VIM-4 remained prominent among Enterobacterales. Strikingly, VIM spread followed distinct host-specific evolutionary strategies. In *Pseudomonas aeruginosa*, dissemination was largely clone-driven, with high-risk lineages ST111 and ST235 supporting long-term persistence through lineage expansion and stable inheritance. By contrast, *Enterobacterales* were dominated by horizontal transmission through broad-host-range IncHI2A, IncA, and IncC plasmids, although only a limited subset of plasmid–VIM combinations achieved intercontinental spread. Among 2,828 loci with sufficient flanking sequence, 96.2% were embedded within integrative genetic elements, frequently nested with insertion sequences and phage-related elements, revealing a multilayered mobile-element network underlying VIM persistence. Structural analyses showed a highly conserved metallo-β-lactamase scaffold but recurrent diversification near substrate-interacting residues, particularly positions 224 and 228. Shared genetic clusters between human-associated and environmental isolates further suggested cross-niche circulation. Together, these findings establish a hierarchical, host-dependent framework for global VIM dissemination, integrating clonal expansion, plasmid transfer, and nested mobile genetic elements.

**Importance:** By analysing 5,617 *bla*_VIM_-positive genomes worldwide, this study reveals a host-dependent transmission architecture of VIM carbapenemases. In *P. aeruginosa*, VIM dissemination is mainly driven by expansion of successful high-risk clones, particularly ST111 and ST235, whereas in *Enterobacterales* it is primarily mediated by broad-host-range plasmids enabling cross-species transfer. Global VIM spread is not driven by a single dominant pathway but by a limited number of successful clone-plasmid-mobile element combinations, while most genetic backgrounds remain regionally restricted. These findings define a hierarchical transmission model of VIM evolution and provide insights for genomic surveillance and targeted control of metallo-β-lactamase-mediated antimicrobial resistance.

## Introduction

The global dissemination of carbapenemase-producing Gram-negative bacteria represents a major threat to modern healthcare. The World Health Organization lists carbapenem-resistant Gram-negative pathogens among the highest-priority bacterial threats for which new therapeutic and control strategies are urgently needed1^1^. Among the major carbapenemase families, KPC, NDM, OXA, IMP, and Verona integron-encoded metallo-β-lactamase (VIM) are particularly important because of their widespread distribution and ability to compromise carbapenem therapy^2^. Since the first description of VIM in *Pseudomonas aeruginosa* in Verona, Italy, in 19973, VIM-producing organisms have been reported across diverse bacterial species and geographic regions^3^. VIM enzymes hydrolyse most β-lactams, with the notable exception of monobactams, and their zinc-dependent catalytic mechanism renders most conventional β-lactamase inhibitors ineffective. Consequently, VIM-producing bacteria remain clinically difficult to treat and epidemiologically challenging to control.^4, 5^.

To date, 96 VIM variants have been registered in the BLDB database (http://www.bldb.eu/), reflecting a complex and continually expanding molecular family^6^. On the basis of protein sequence homology, VIM enzymes can be broadly grouped into three major sublineages represented by VIM-1, VIM-2, and the highly divergent VIM-7^7^. These sublineages differ in host distribution. VIM-2 is the most widely disseminated variant globally and is carried predominantly by *P. aeruginosa*, whereas VIM-1 is more frequently associated with *Enterobacterales*^8, 9^. Such host preference may reflect differences in signal peptide processing, expression efficiency, and fitness costs across bacterial backgrounds^10^. Despite this sequence diversity, the core catalytic scaffold of VIM enzymes is highly conserved, substitutions at key residues, including positions 224 and 228, can alter substrate profiles or catalytic kinetics^11–14^, suggesting that VIM diversification occurs within a strongly constrained functional framework.

The global persistence of *bla*_VIM_ is closely linked to its association with multiple layers of mobile genetic elements. *bla*_VIM_ is typically embedded as a gene cassette within class 1 integrons^15, 16^, which can nested within larger mobile genetic elements, and other mobile genetic structures^17–21^. Broad-host-range plasmids, including IncA/C- and IncHI2-related backbones, can facilitate horizontal dissemination across bacterial species, whereas chromosomal integration may promote stable maintenance within successful clonal lineages^22–25^. Thus, *bla*_VIM_ transmission is unlikely to depend on a single genetic vehicle. Rather, its long-term dissemination may reflect interactions among bacterial host background, clonal expansion, plasmid transfer, and nested mobile genetic elements.

Ecologically, VIM dissemination has extended beyond clinical isolates and shows clear evidence of environmental persistence. Although VIM was initially associated with hospital outbreaks in Southern Europe^26–28^, accumulating evidence identifies hospital water systems, including sinks and drains, as important secondary reservoirs^29–31^. Multi-year investigations have shown that persistent environmental colonisation can drive recurrent outbreaks of VIM-positive strains, rather than direct patient-to-patient transmission alone^32–34^. This patient-environment-patient cycle makes VIM circulation particularly difficult to interrupt within a One Health framework.

Therapeutically, ceftazidime-avibactam is widely used for infections caused by KPC-producing carbapenem-resistant Gram-negative bacteria^35^. In contrast, VIM-producing organisms are intrinsically resistant to ceftazidime-avibactam, and clinicians often rely on cefiderocol or aztreonam-avibactam as salvage options^36^. The increasing use of these agents may reshape the resistance landscape under antibiotic selection. Cefiderocol resistance has already been linked to mutations affecting siderophore uptake, whereas intrinsic resistance to ceftazidime-avibactam may favour the persistence and diversification of MBL-producing strains. VIM subtypes such as VIM-2, VIM-4, and VIM-11 differ in hydrolytic activity, genetic background, and transmissibility. Some variants may compromise salvage regimens and create new clinical management challenges^37–39^.

Despite extensive regional studies over the past two decades, the global transmission architecture of VIM remain incompletely understood. Previous studies have highlighted the roles of high-risk clones, transferable plasmids, and mobile genetic elements, but these components have largely been examined separately or at regional scales. It therefore remains unclear whether global VIM dissemination is driven by common transmission platforms or distinct host-dependent strategies, and why only certain VIM–host–vector combinations achieve widespread geographic dissemination. Here, we integrate publicly available *bla*_VIM_-positive genomes collected worldwide from 1999 to 2025 to define the global molecular epidemiology of this resistance determinant. We characterise the spatiotemporal trajectories of major *bla*_VIM_ subtypes and dissect their transmission through clonal expansion, plasmid-vector turnover, and hierarchical mobile-element integration. This large-scale genomic analysis provides a global framework for understanding *bla*_VIM_ evolution and transmission. It also supports One Health-informed strategies to control MBL-mediated carbapenem resistance.

## Methods

### Genome acquisition, species identification, and genotyping

Genome assemblies carrying *bla*_VIM_ variants were retrieved from the NCBI Pathogen Isolate Browser using the search query AMR_genotypes:blaVIM*^40^. The search covered isolates from multiple BioProjects, and 6,697 assembled genomes were downloaded directly from the Pathogen Isolate Browser. After merging, de-duplication, and quality filtering, 5,617 high-quality assemblies were retained for downstream comparative genomic analyses^41^.

Sample metadata were extracted from the corresponding NCBI BioSample records. For isolates without explicit collection dates, and for BioProjects contributing fewer than 50 isolates, the sequence submission date was used as an approximate temporal proxy. Assembly quality was assessed with CheckM2. Genomes were included only if they had an N50 of at least 20,000 bp and no more than 500 contigs (Supplementary Data 1)^42, 43^.

Species-level taxonomic assignment was performed using Speciator. Prokka v1.14.6 was used for rapid prokaryotic genome annotation, and antimicrobial resistance genes were identified and classified using AMRFinderPlus v3.12.8^44, 45^. Only *bla*_VIM_ alleles with 100% nucleotide identity and 100% query coverage were considered valid VIM variants (Supplementary Data 1).

In silico multilocus sequence typing (MLST) was performed using mlst v2.19.0^46^. Several *bla*_VIM_-carrying species lack standardised or fully curated MLST schemes, including *Morganella morganii*, *Proteus mirabilis*, *Providencia rettgeri*, *Providencia stuartii*, *Serratia marcescens*, and *Stenotrophomonas maltophilia*. Therefore, a two-step hierarchical lineage-classification pipeline was implemented for these taxa. First, untyped isolates were assigned to species-level groups using MASH v2.3 with a distance threshold of ≤0.05^47^. PopPUNK v4.2.0 was then applied within each MASH-defined group to infer intra-species lineages^48^. PopPUNK databases were constructed with --sketch-size 1000000, --min-k 15, --max-k 29, and --qc-filter prune. Taxonomic inspection revealed extensive genetic divergence within *Providencia rettgeri* and *Providencia stuartii*. Five and two distinct subgroups, respectively, exceeded the conventional 0.05 MASH distance threshold. Intra-group MASH distances reached 0.19, suggesting that these groups may represent cryptic species or unrecognised genera^48^. Because formal taxonomic revision was beyond the scope of this study, these lineages were provisionally designated as *Providencia rettgeri* CG1, *Providencia rettgeri* CG2, and so forth.

For assemblies lacking explicit sequencing-platform information, a heuristic classification rule was applied. Genomes with fewer than 20 contigs were classified as long-read or hybrid assemblies, whereas those with 20 or more contigs were classified as short-read assemblies. Countries were assigned to regions and subregions according to the United Nations geoscheme.

### Assembly dereplication and VIM-cluster definition

To reduce bias introduced by clonal outbreaks, we generated a dereplicated dataset by selecting representative genomes from genetically similar isolates. Pairwise core-genome single-nucleotide variant (SNV) distances within identical STs or clonal groups were calculated using the ska fasta and ska distance commands in SKA v1.0.0^49^. Genomes were considered closely related when they differed by ≤5 SNVs per megabase pair. Graph-based clustering was then performed in igraph v2.0.3 using SNV distances as edge weights^50^. VIM clusters were defined as genome groups that carried the same VIM variant, belonged to the same species, shared the same ST or clonal group, and differed by ≤5 SNVs per megabase pair.

### Plasmid analysis

Plasmid replicon typing and contig binning were performed using mob_recon from MOB-suite v3.1.8, which assigns contigs to plasmids and infers incompatibility types^51^. Plasmid analyses were restricted to long-read or hybrid assemblies, yielding 224 plasmid-derived contigs. A custom *bla*_VIM_-positive plasmid database was built from these contigs using mob_typer --multi followed by mob_cluster --mode build. Short-read assemblies were then screened against this custom database using mob_recon to identify plasmid-homology signals.

### Genetic context and mobile genetic element analysis

The 10-kb genomic regions upstream and downstream of *bla*_VIM_ were extracted and clustered using Flanker v0.1.5 with the parameters --flank both --window 10000 --gene blaVIM --include_gene --cluster^52^. Only contig segments of at least 6 kb were retained for mobile genetic element annotation. Mobile genetic elements within the extracted regions were identified by sequence similarity searches against mobileOG-db. Putative mobility-related genes were annotated according to the mobileOG-db classification system, including insertion sequence or transposase-associated genes, integrase or recombinase genes, plasmid-associated genes, phage-related genes, and other mobility-related elements. The distribution and organisation of these features were then examined to characterise the genetic environments surrounding *bla*_VIM_^53–55^.

### Protein sequence and structural analysis

Signal peptide cleavage sites in VIM protein sequences were predicted using SignalP v6.0^56^. Full-length mature VIM sequences were aligned with Clustal Omega v1.2.4^57^. Phylogenetic trees inferred from multiple sequence alignments were annotated and visualised using ggtree v3.16.3. Amino acid conservation scores were calculated from the alignment using the conserve function in the bio3d v2.4-4 R package with method = “similarity” and sub.matrix = “bio3d”^58^. Three-dimensional structural models of mature VIM enzymes were predicted using ColabFold v1.5.5 with the AlphaFold2 algorithm, and the top-ranked models were retained for analysis^59, 60^. Per-residue structural conservation scores were calculated using FoldMason v1.763a428^61^.

### Statistical analysis and visualization

All statistical analyses and visualizations were performed in R v4.4.1 with the following R packages: tidyverse v2.0.0, colorspace v2.1-1, viridis v0.6.5, ggh4x v0.2.8, ggstream v0.1.0, maps v3.4.2, scatterpie v0.2.4, sf v1.0-17, rnaturalearth v1.0.1, ggnewscale v0.5.0, treemapify v2.5.6, patchwork v1.3.0, igraph v2.0.3, qgraph v1.9.8, ggraph v2.2.1, ggforce v0.5.0, ggalluvial v0.12.5, ggtree v3.12.0, treeio v1.28.0, and aplot v0.2.3^46, 57, 62–68^.

## Results

### 1. Global transition of *bla*_VIM_ epidemiology reveals dominance of a limited number of successful variants

To characterise the global epidemiology of *bla*_VIM_-positive strains, we analysed 5,617 genomes collected primarily between 1999 and 2025, including one historical record from 1905. The dataset covered 73 countries or regions across six continents. We identified 40 *bla*_VIM_ subtypes across 16 bacterial genera, revealing substantial allelic diversity and a broad host range.

The geographic distribution of *bla*_VIM_ subtypes was highly heterogeneous (Figure 1A). *bla*_VIM-2_ and *bla*_VIM-1_ were the dominant globally distributed subtypes. Both were prevalent across Europe, Asia, and the Americas, with major epidemic centres in North America and Europe. By contrast, *bla*_VIM-4_ had a more restricted distribution and was concentrated mainly in Europe, with sporadic detection in Asia and South America. Other low-frequency subtypes, including *bla*_VIM-5_, *bla*_VIM-20_, and additional rare VIM variants, were scattered and did not form clear regional epidemics.

**Figure 1.**
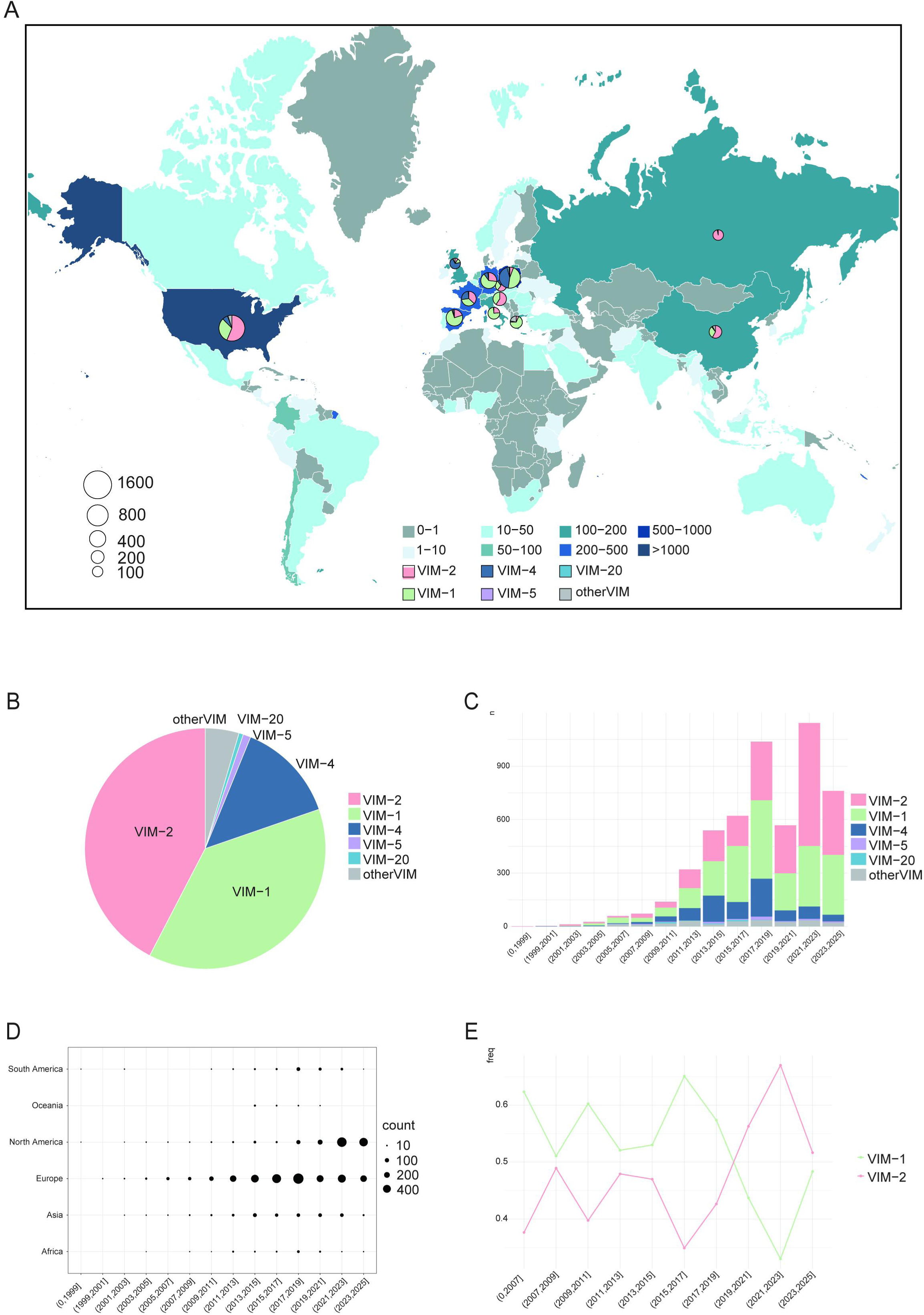
Geographic and temporal dissemination of VIM carbapenemase genes. (A) Global map showing the geographic distribution and species composition of VIM-carrying genomes; pie-chart size corresponds to isolate number. VIM variants represented by fewer than 100 isolates are grouped as “Other VIM”. (B and C) Temporal prevalence dynamics of dominant VIM variants worldwide. (D) Dot plot showing yearly counts of VIM-positive genomes. (E) Prevalence trends of VIM-1 and VIM-2. Source data are provided in the Source Data file.

Overall, *bla*_VIM-2_ was the most common subtype, accounting for 42.38% of isolates (2,381/5,617). It was followed by *bla*_VIM-1_ (37.90%, 2,129/5,617) and *bla*_VIM-4_ (13.55%, 761/5,617). Together, these three subtypes represented more than 90% of all isolates, whereas the remaining variants were individually rare (Figure 1B). These findings indicate that the global *bla*_VIM_ burden is driven primarily by a limited number of dominant variants (Supplementary Data 1).

Temporal analysis showed an overall increase in detected *bla*_VIM_-positive strains during the study period, with apparent slowdowns during 2019–2021 and 2023–2025 (Figure 1C). *bla*_VIM-1_ and *bla*_VIM-2_ remained the dominant subtypes throughout the study period but showed a clear dynamic shift. Before 2019, *bla*_VIM-1_ was slightly more prevalent. Thereafter, *bla*_VIM-2_ increased and gradually became the dominant globally circulating subtype, indicating a transition from *bla*_VIM-1_-dominated to *bla*_VIM-2_-dominated epidemiology (Figures 1C and 1E).

Subtype-specific temporal patterns were also observed. *bla*_VIM-1_ declined transiently during 2019–2021 and then stabilised, whereas *bla*_VIM-2_ plateaued during both 2019–2021 and 2023–2025. *bla*_VIM-4_ and other VIM variants increased before 2015 but gradually decreased thereafter. *bla*_VIM-5_ peaked briefly during 2017–2019, while *bla*_VIM-20_ was detected mainly between 2009 and 2017 and became rare thereafter (Figure 1C).

### 2. Host-specific transmission strategies shape global VIM dissemination

We next analysed the host distribution of the major epidemic subtypes *bla*_VIM-1_, *bla*_VIM-2_, and *bla*_VIM-4_. Overall, *bla*_VIM_ was distributed across multiple Gram-negative bacilli, indicating substantial cross-genus transmissibility. Each subtype, however, showed a distinct host preference (Figures 2A–F and Supplementary Data 2).

**Figure 2.**
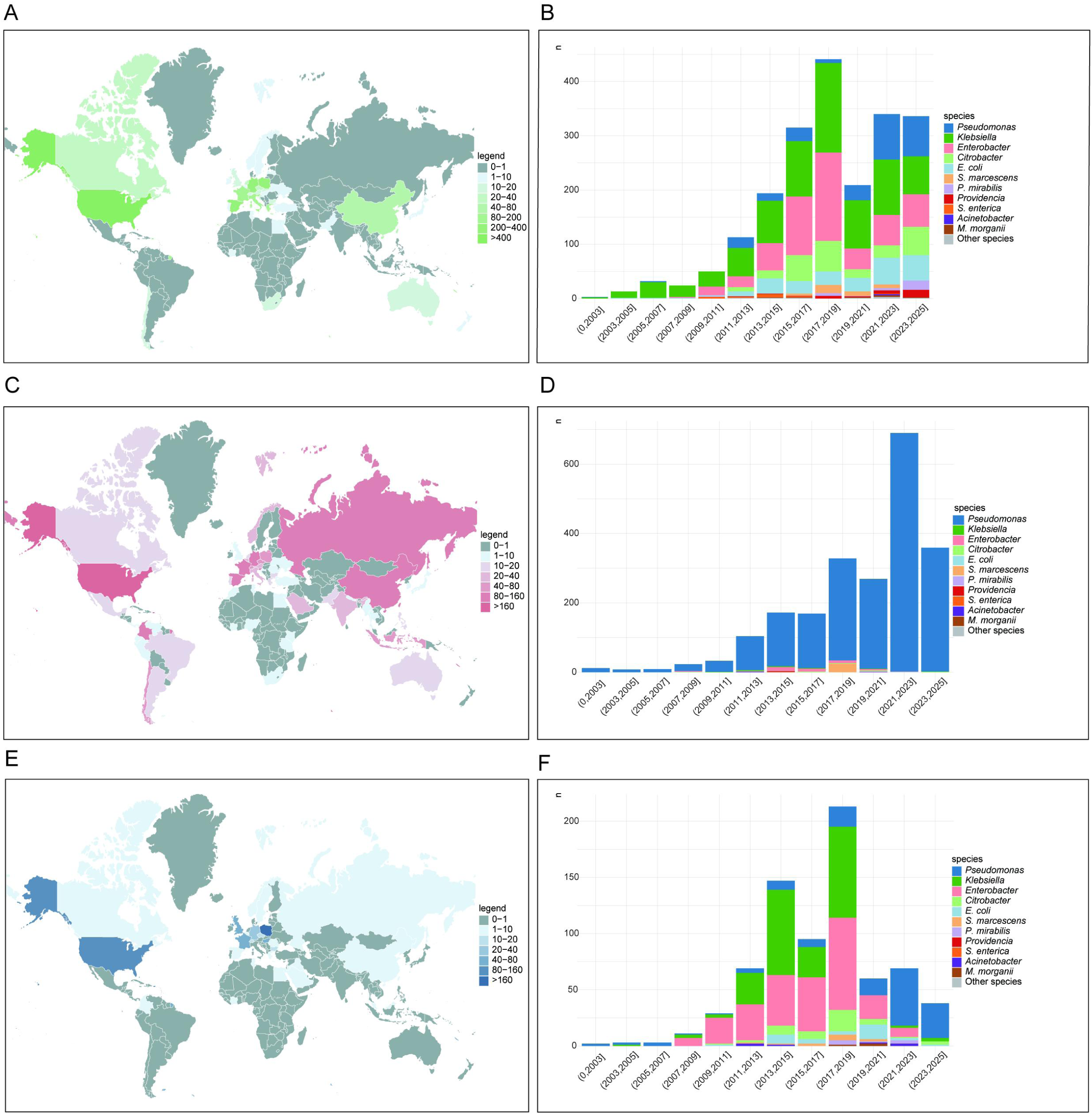
Epidemiology of VIM-1, VIM-2, and VIM-4. (A) Global distribution and regional endemic patterns of VIM-2. (B) Temporal changes in species composition among VIM-2 genomes. (C) Global distribution and regional endemic patterns of VIM-1. (D) Temporal changes in species composition among VIM-1 genomes. (E) Global distribution and regional endemic patterns of VIM-4. (F) Temporal changes in species composition among VIM-4 genomes.

#### 2.1 *P. aeruginosa* : clone-driven dissemination

At the genus level, *P. aeruginosa* was the predominant host and represented the major reservoir of *bla*_VIM-2_-positive isolates, suggesting a strong association between this subtype and specific bacterial lineages. *Klebsiella* and *Enterobacter* formed the second major host group and were highly represented among *bla*_VIM-1_- and *bla*_VIM-4_-positive isolates. *Citrobacter*, *Escherichia coli*, *Serratia marcescens*, and *Acinetobacter* were also detected at appreciable frequencies, indicating broad host adaptability of *bla*_VIM_.

Host distributions differed markedly among subtypes. *bla*_VIM-2_ was strongly concentrated in *P. aeruginosa*, with only limited detection in *Enterobacter* and *S. marcescens* (Figures 2A and 2B). This pattern indicated pronounced host specificity. By contrast, *bla*_VIM-1_ and *bla*_VIM-4_ had broader host ranges and occurred in multiple *Enterobacterales* and non-fermenting Gram-negative bacteria, suggesting higher cross-species transmission potential (Figures 2C–F). Together, these findings indicate that *bla*_VIM-2_ dissemination in *P. aeruginosa* is characterised by strong host specificity and preferential association with successful bacterial lineages, consistent with a clone-associated persistence strategy.

#### 2.2 *Enterobacterales*: plasmid-driven dissemination

In contrast to the lineage-associated pattern observed in *P. aeruginosa*, *bla*_VIM-1_ and *bla*_VIM-4_ showed broader host distributions among *Enterobacterales*, suggesting a greater contribution from horizontal dissemination. Temporal dynamics further revealed shifts in host structure. For *bla*_VIM-1_, *Klebsiella* and *Enterobacter* dominated and increased from 2003 to 2019. After 2019, their relative contribution declined, whereas the proportions of *P. aeruginosa* and *Klebsiella* increased (Figure 2D). The host structure of *bla*_VIM-2_ remained comparatively stable, with *P. aeruginosa* dominant throughout and gradually increasing in proportion (Figure 2B). The pattern for *bla*_VIM-4_ resembled that of *bla*_VIM-1_. *Klebsiella* and *Enterobacter* dominated early and increased from 2003 to 2019, followed by an overall decline after 2019 and a relative increase in *P. aeruginosa*, which surpassed *Enterobacter* as the main host after 2021 (Figure 2F).

### 3. Successful high-risk clones act as major reservoirs for *bla*_VIM_ persistence

After defining global distribution patterns, we examined whether specific bacterial lineages contributed disproportionately to blaVIM persistence. Overall, *bla*_VIM_ was detected across 41 bacterial species in 16 genera, but showed strong associations between specific variants and and particular host lineages (Figure 3A).

**Figure 3.**
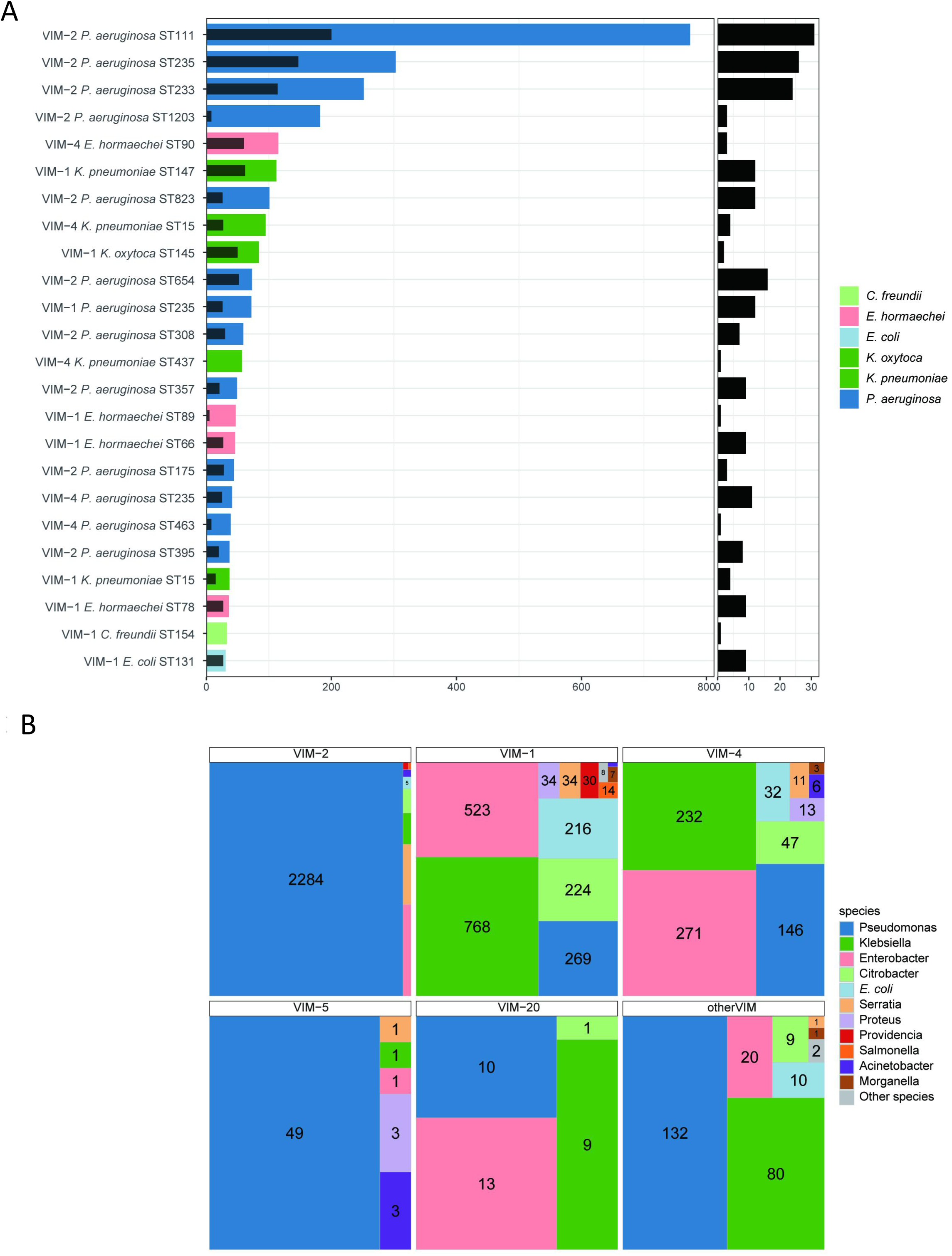
Host species and dominant lineages associated with blaVIM-carrying genomes. (A) Species distribution across distinct *bla*_VIM_ variants. (B) Dominant lineages before and after dereplication, with grey overlays indicating dereplicated genome counts. Lineages containing ≥30 genomes are shown, together with the geographic subregion distribution of each lineage. Detailed information is provided in the supplementary datasets.

At the genus level, *Pseudomonas* was the dominant host group, with 2,890 isolates, approximately 99% of which were *P. aeruginosa*. This species also harboured the greatest diversity of *bla*_VIM_ subtypes. Notably, 95% of *bla*_VIM-2_-positive isolates belonged to *P. aeruginosa* (Figure 3B). *bla*_VIM-5_ and several rare VIM variants were also mainly detected in *Pseudomonas*. By contrast, *Klebsiella* predominantly carried *bla*_VIM-1_, whereas *Enterobacter hormaechei* mainly carried *bla*_VIM-1_, *bla*_VIM-4_, and *bla*_VIM-20_ (Figure 3B).

At the clonal level, several high-prevalence *P. aeruginosa* STs dominated *bla*_VIM_ transmission. ST111 was the most common clone (27.8%, 796 isolates), followed by ST235 (15.4%, 443 isolates) and ST233 (9.4%, 270 isolates) (Figure 3A). These dominant clones carried multiple *bla*_VIM_ subtypes, with ST235 showing the greatest subtype diversity (13 subtypes). Notably, *bla*_VIM-20_ was absent from these major clones and was concentrated in ST175, suggesting reliance on a distinct clonal background (Supplementary Data 1).

Further analysis showed that *bla*_VIM-2_ was overwhelmingly dominant among major *P. aeruginosa* clones, demonstrating a strong coupling between this variant and globally successful epidemic lineages. Beyond *Pseudomonas*, *bla*_VIM-1_ in *Klebsiella* was mainly associated with ST147 and ST15, whereas *bla*_VIM-4_ in *E. hormaechei* was concentrated in ST90. *E. coli* also represented an important host for *bla*_VIM-1_, with ST89 as the representative lineage (Figure 3A). These findings support a model in which successful *P. aeruginosa* clones serve as long-term reservoirs that maintain and amplify *bla*_VIM_ dissemination.

### 4. Broad-host-range plasmids drive cross-species dissemination of *bla*_VIM_ in *Enterobacterales*

To determine whether horizontal transfer contributed to *bla*_VIM_ dissemination, we characterised plasmid localisation and replicon diversity using genomes with long-read assembly data. In the long-read dataset (n = 349), *bla*_VIM_ was predominantly plasmid associated. Overall, 58.45% of genomes (204/349) carried *bla*_VIM_ on plasmid-derived contigs. MOB-suite clustering assigned these *bla*_VIM_-positive plasmids to 38 clusters carrying nine distinct *bla*_VIM_ variants, indicating dissemination across multiple plasmid backgrounds.

IncHI2A, IncA, IncC, IncL/M, and IncFIB were the major plasmid types carrying *bla*_VIM_. IncHI2A, IncA, and IncC represented primary plasmid backbones, each corresponding to large plasmid clusters distributed across multiple countries and host species (Figures 4A, 4B and Table 1). Our analysis revealed that *bla*_VIM_-associated plasmids circulated across a broad bacterial host spectrum, with IncA/IncC plasmids detected across 22 species, supporting their role as important cross-species transmission platforms. IncA and IncC had broad host ranges spanning 22 species, including *Klebsiella pneumoniae*, *E. hormaechei*, and *Citrobacter freundii*. They mainly carried *bla*_VIM-1_ and *bla*_VIM-4_ and served as key vectors for global *bla*_VIM-1_ dissemination. IncHI2A also crossed host-species and national boundaries, indicating broad dissemination capacity. By contrast, IncL/M and IncFIB had narrower host ranges, with *K. pneumoniae* as a major host, and predominantly carried *bla*_VIM-1_ (Figure 4A).

**Figure 4.**
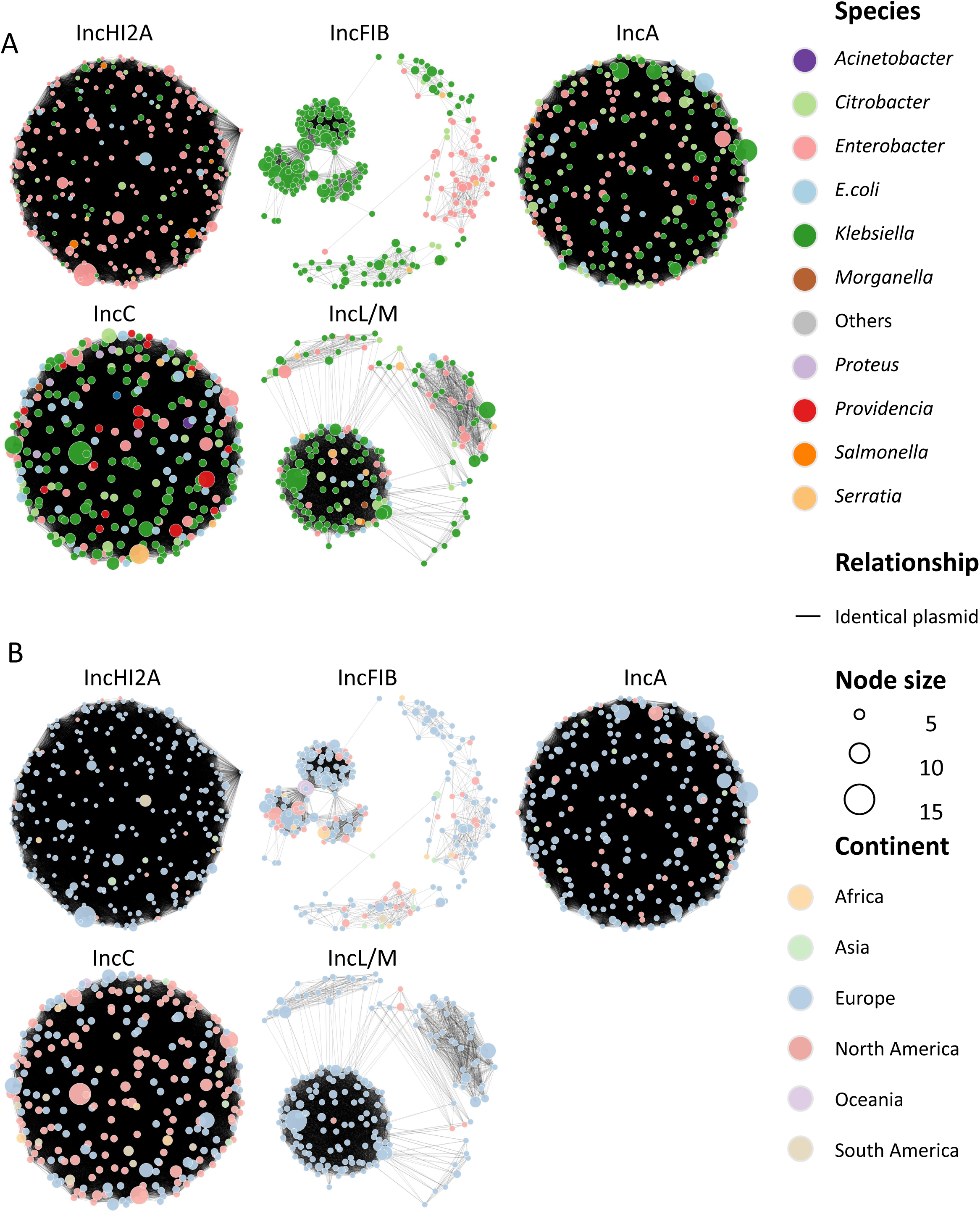
Major *bla*_VIM_-harboring plasmids across bacterial species and geographic regions. (A) Plasmid network grouped by incompatibility type and coloured by host bacterial species. (B) The same plasmid network coloured by geographic region. Nodes represent VIM clusters, and edges indicate shared identical plasmids among clusters. Node size corresponds to the number of genomes and countries carrying the same plasmid.

**Table 1.**
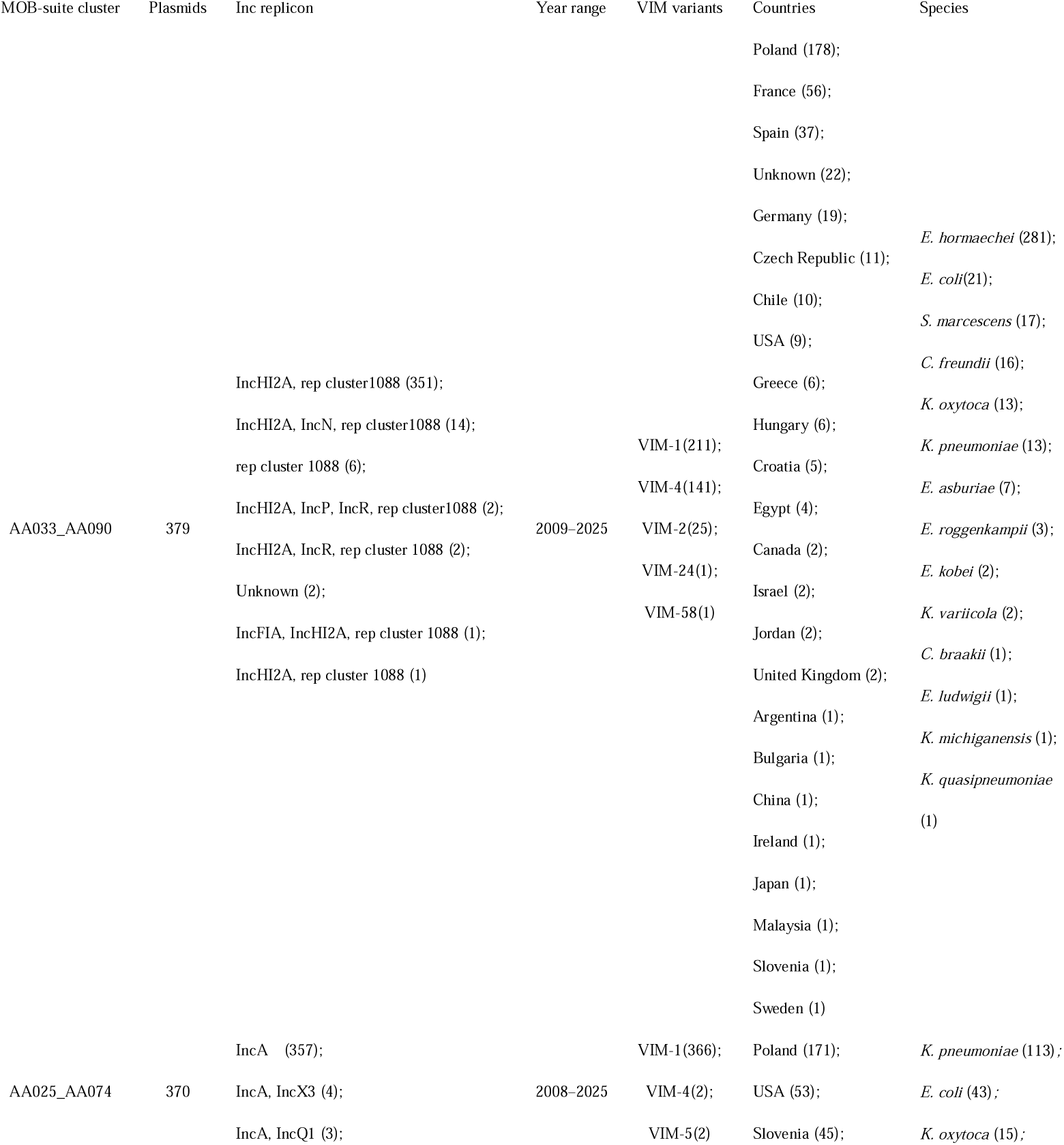
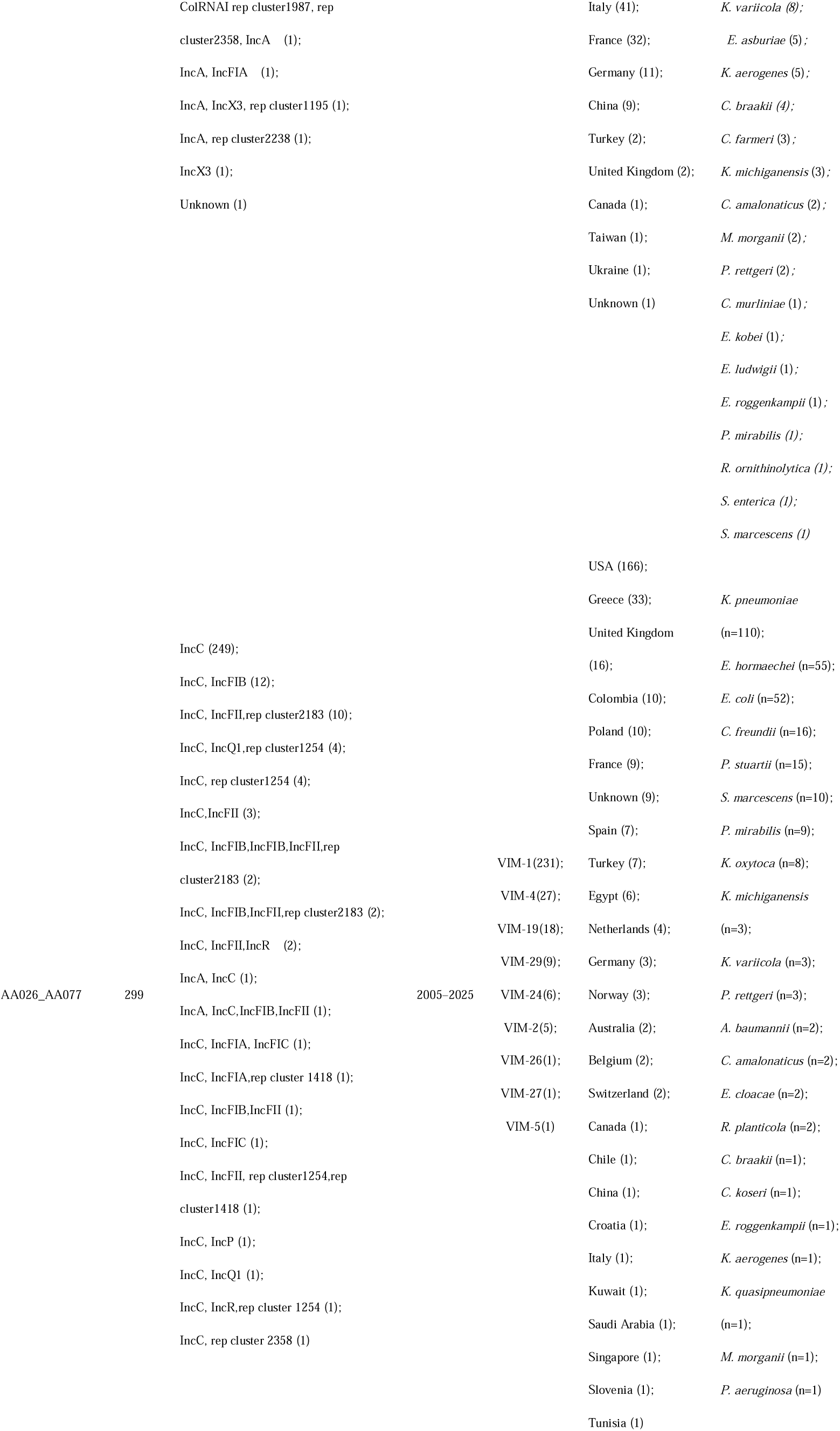
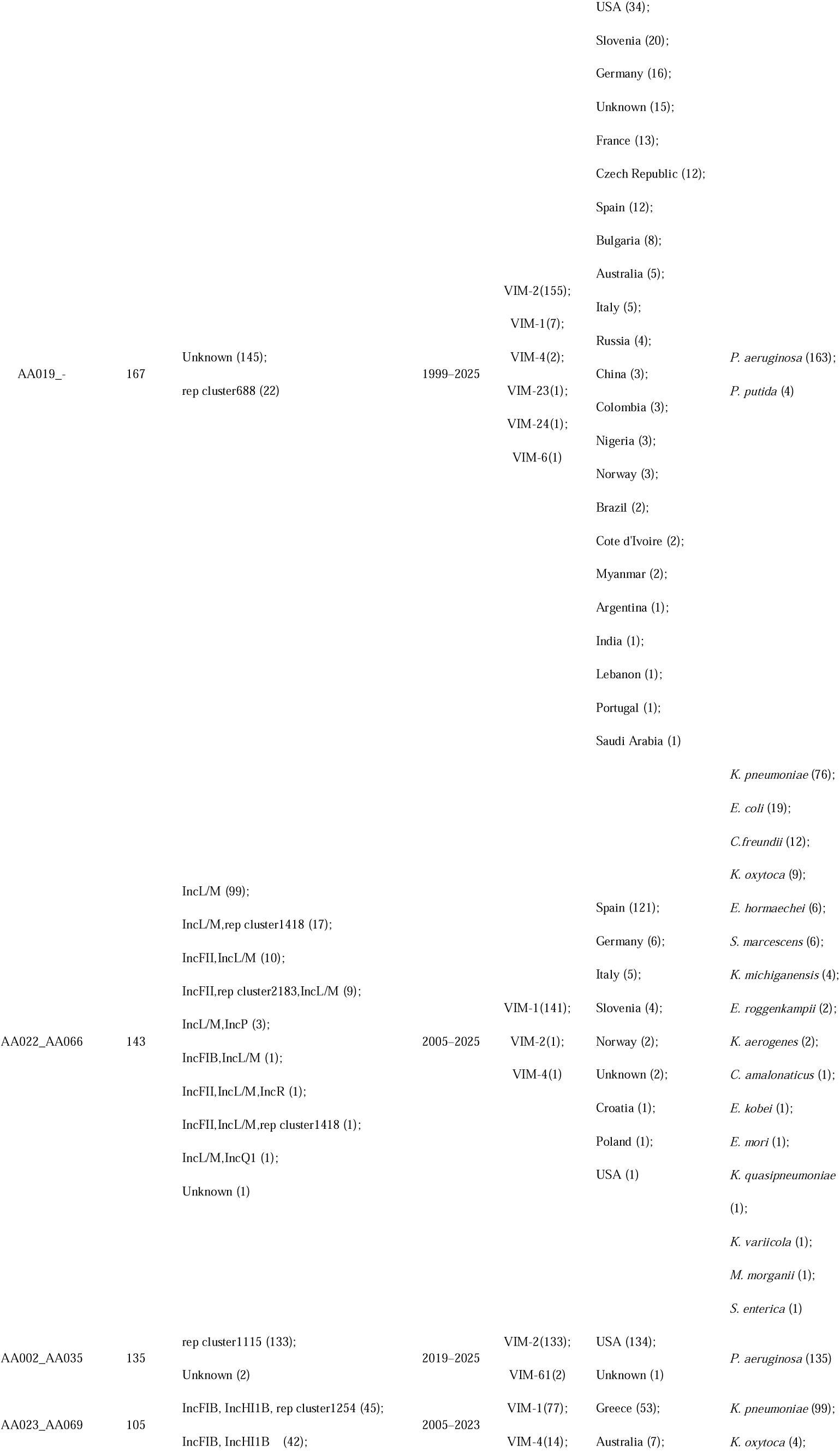
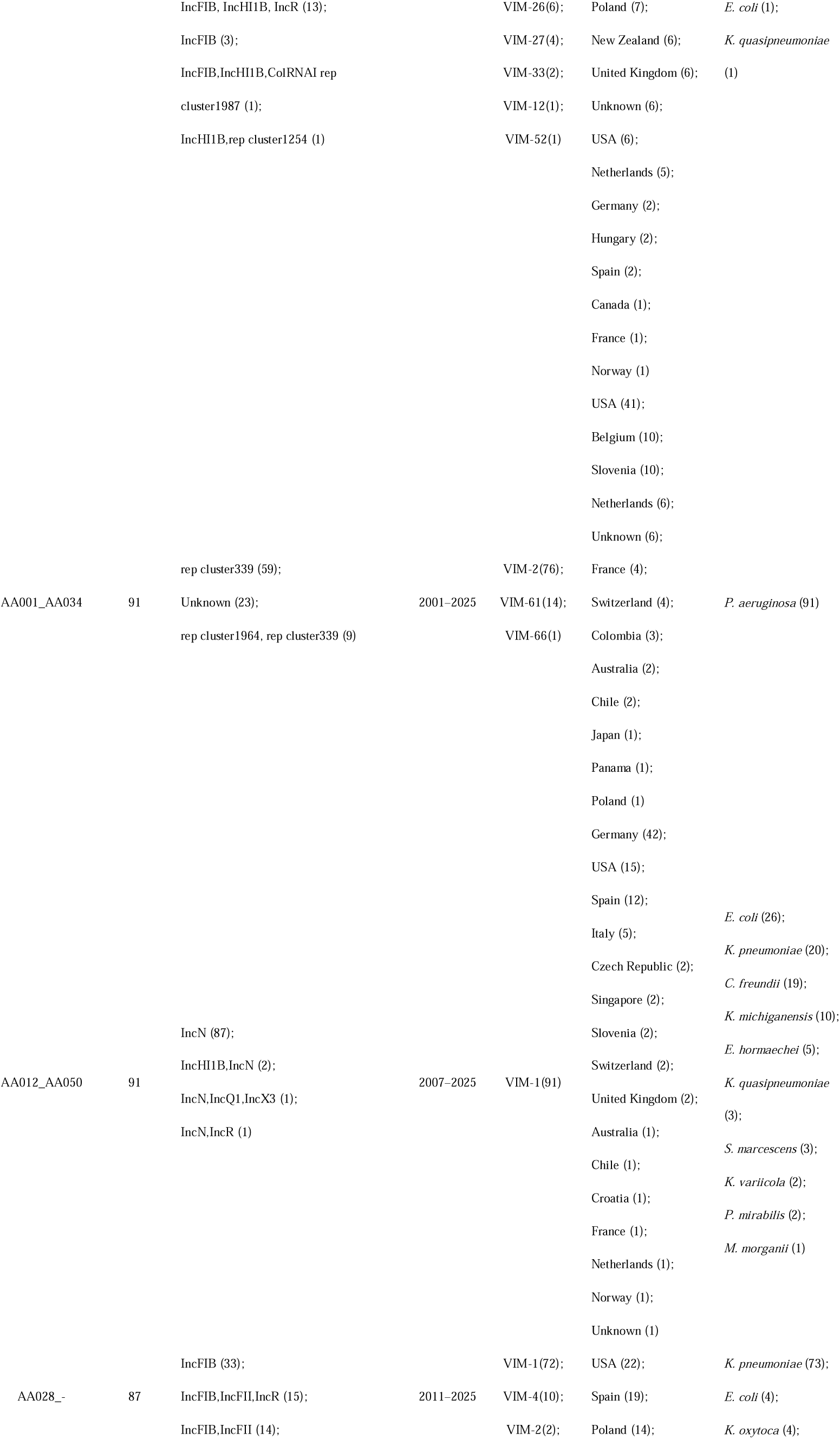
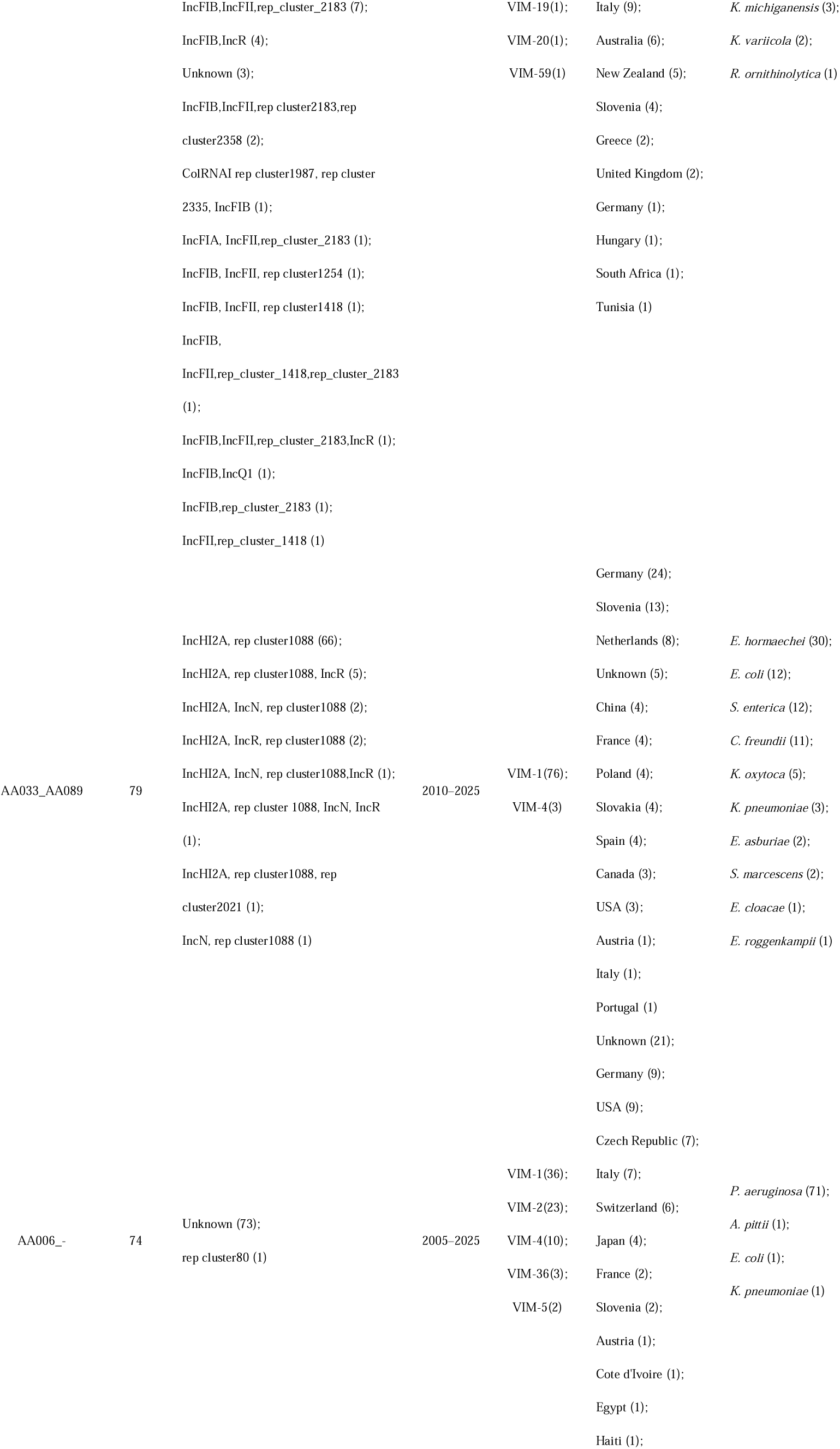
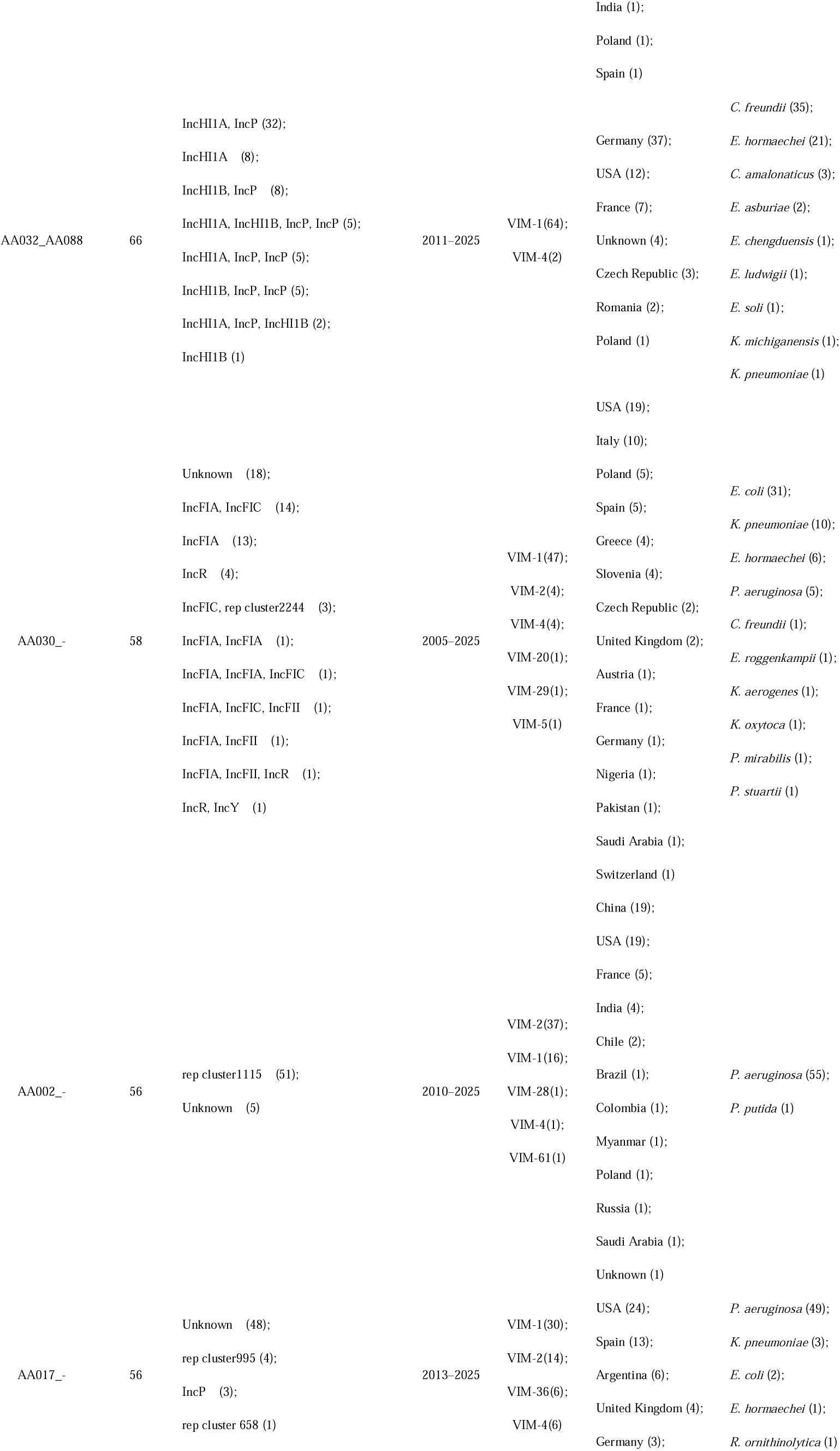
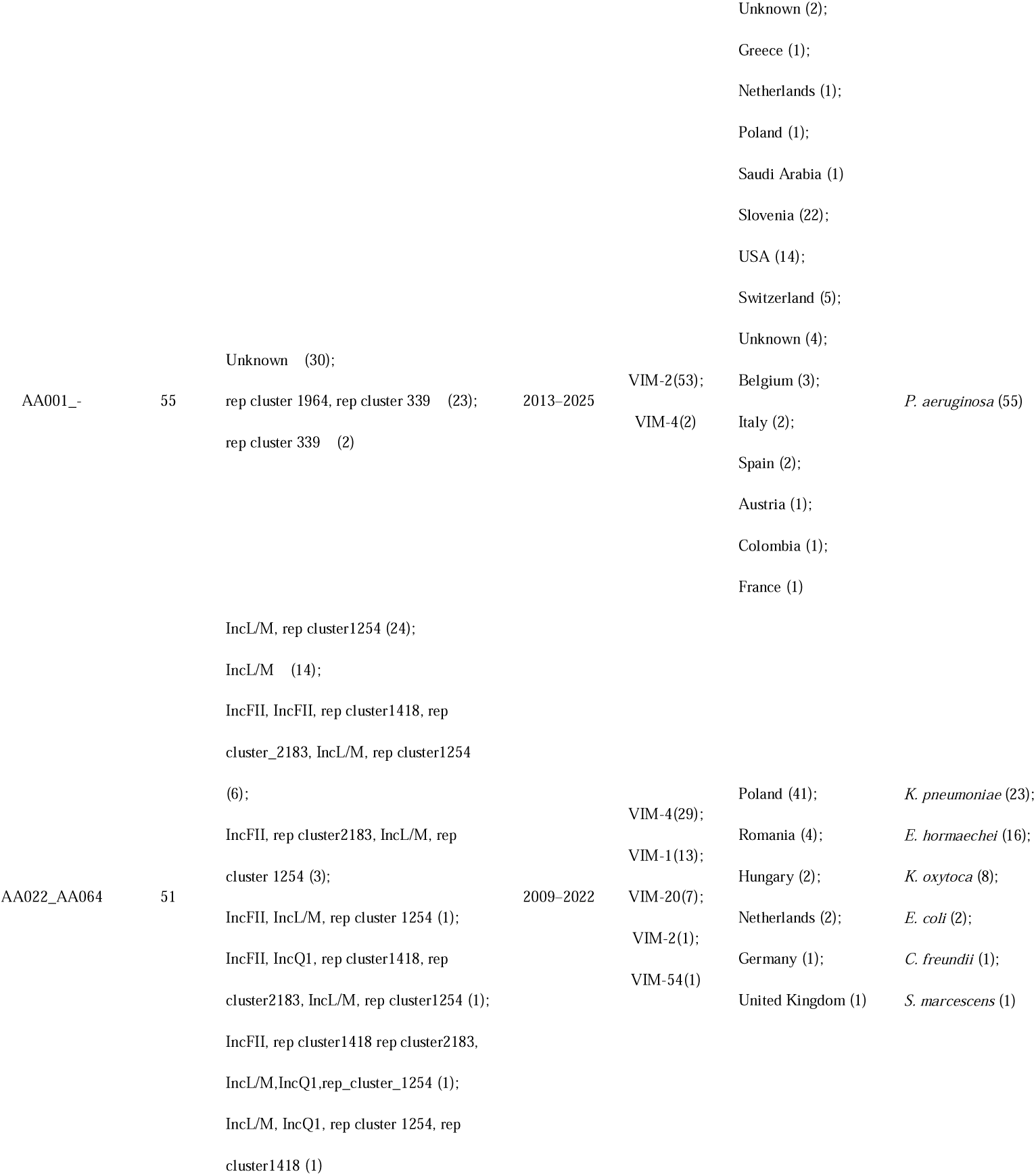
Summary of key plasmid clusters (n ≥ 50) and associated features.

Geographically, *bla*_VIM_-positive plasmids displayed a highly heterogeneous dissemination pattern: a small number of plasmid–variant combinations achieved cross-country or intercontinental spread, whereas most remained restricted to individual countries or regions. (Figure 4B). IncHI2A, IncA, IncL/M, and IncFIB plasmids were concentrated in Europe. IncC plasmids were detected predominantly in North America, especially the United States. Among the 38 VIM variant-plasmid cluster combinations, only a small subset showed cross-country or intercontinental spread. Such dissemination was mediated mainly by broad-host-range plasmids, including IncHI2A, IncA, IncC, IncL/M, and IncFIB, which repeatedly moved across host species. Most combinations remained confined to individual countries or regions. Plasmid-sharing network analysis further revealed frequent exchange of *bla*_VIM_-positive plasmids among host species (Figure S3). *Enterobacterales*, represented by *Enterobacter* spp., *Klebsiella* spp., and E. coli, occupied central network positions. These patterns indicate pivotal roles for *Enterobacterales* in horizontal plasmid-mediated transfer of *bla*_VIM_.

The genetic context of *bla*_VIM_ in *P. aeruginosa* differed markedly from that in *Enterobacterales*. In *P. aeruginosa*, *bla*_VIM-2_ was associated mainly with non-canonical or unclassified plasmid backbones, including IncP, rep cluster 339, rep cluster 724, and rep cluster 2338. A high proportion of *bla*_VIM_ loci were chromosomal or otherwise unclassified (Figure 4A). Moreover, *bla*_VIM_ in *P. aeruginosa* showed strong lineage-specific clustering, particularly within the high-risk clone ST235. These contrasting patterns indicate that *bla*_VIM_ dissemination follows distinct host-dependent strategies: plasmid-mediated horizontal expansion predominates among *Enterobacterales*, whereas lineage-specific persistence contributes more substantially in *P. aeruginosa*.

### 5. Nested mobile genetic element architectures facilitate *bla*_VIM_ persistence

To characterise the mobile genetic environment of *bla*_VIM_, we screened the 10-kb regions upstream and downstream of each *bla*_VIM_ locus. Regions of at least 6 kb were retained for analysis, yielding 2,828 eligible *bla*_VIM_-positive regions (Figure 5).

**Figure 5.**
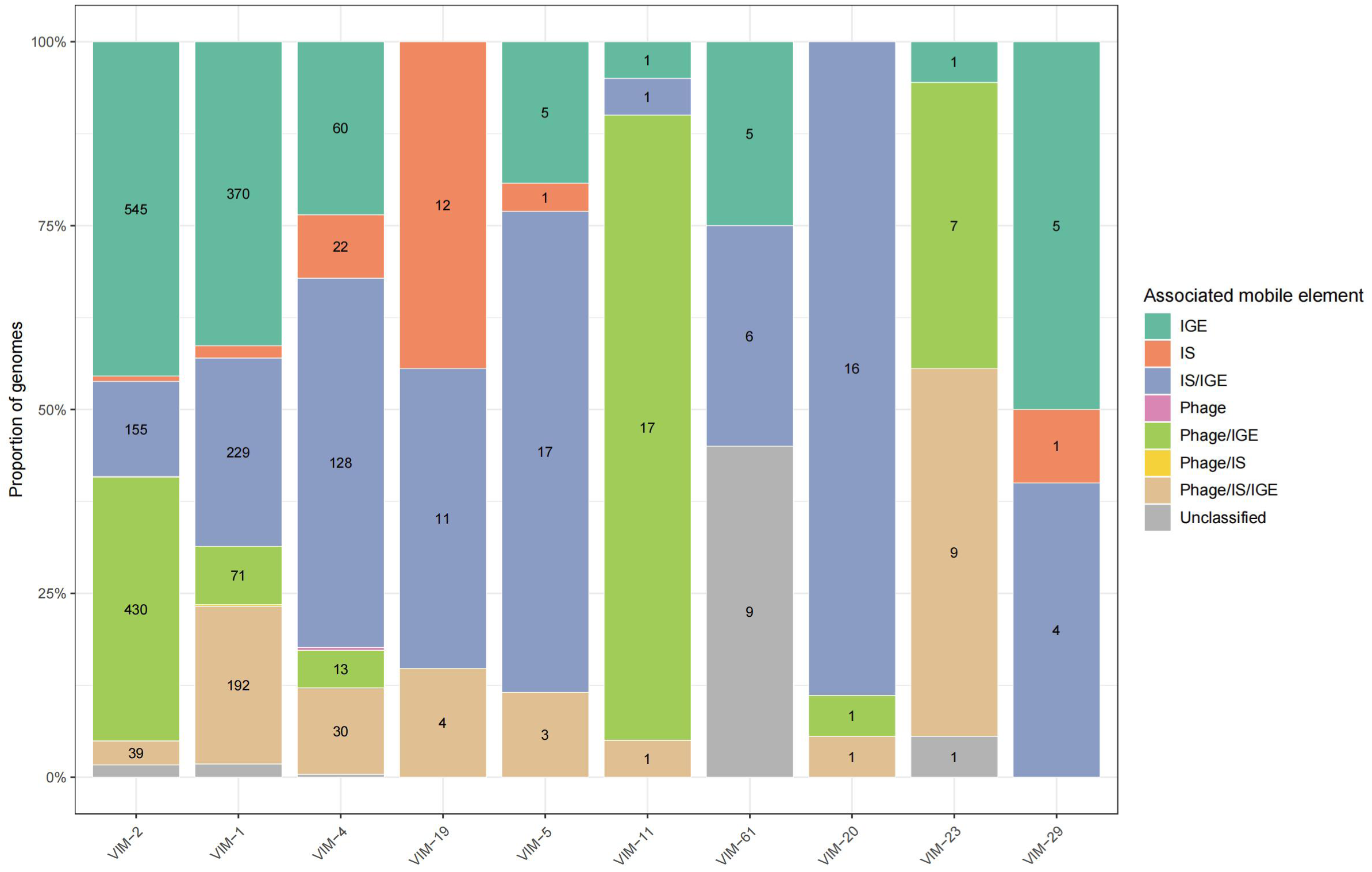
Association of major *bla*_VIM_ variants with mobile genetic elements. Bar chart showing the proportion of each *bla*_VIM_ variant associated with different mobile genetic elements, including integrative genetic elements, insertion sequences, phage-related elements, integrons, and unclassified contexts. Numerical values for each category are labelled within the corresponding bars. Raw data are provided in Supplementary Data 1.

Overall, *bla*_VIM_ was strongly associated with mobile genetic elements. In total, 96.2% of *bla*_VIM_-positive regions were linked to integrative genetic elements, with integrative genetic elements alone accounting for 40.2%. More frequently, these elements co-occurred with other mobile elements, including integrative genetic element/insertion sequence (23.0%), integrative genetic element/phage (21.8%), and integrative genetic element/insertion sequence/phage combinations (11.3%) (Figure 5). These results indicate that the genetic context of *bla*_VIM_ is centred on integrative genetic elements and often forms composite structures with other mobile elements. These findings indicate that *bla*_VIM_ dissemination is supported by multilayered genetic platforms rather than by a single mobile element type.

The major epidemic subtypes *bla*_VIM-2_ and *bla*_VIM-1_ both showed integrative genetic elements as the predominant associated category (45.5% and 41.3%, respectively). Their co-localisation patterns differed, however. *bla*_VIM-2_ was more frequently linked to phage/integrative genetic element contexts, whereas *bla*_VIM-1_ was more often associated with insertion sequence/integrative genetic element contexts (Figure 5).

Several low-frequency subtypes displayed more specific mobile-element association patterns. For example, *bla*_VIM-11_ was mainly co-localised with phage/integrative genetic element contexts (85.0%), whereas *bla*_VIM-20_ was dominated by insertion sequence/integrative genetic element contexts (88.9%). The triple co-occurrence of phage, insertion sequences, and integrative genetic elements was relatively frequent for *bla*_VIM-23_ (50.0%). By contrast, *bla*_VIM-19_ was more commonly associated with insertion sequences alone, and *bla*_VIM-61_ showed a high proportion of unclassified contexts (45.0%) (Figure 5).

### 6. Sequence and structural analysis of VIM variants

To clarify the evolutionary relationships and potential structural divergence among VIM variants, we integrated phylogenetic, sequence-conservation, and protein-structure analyses (Figure 6 and Supplementary Data 3).

**Figure 6.**
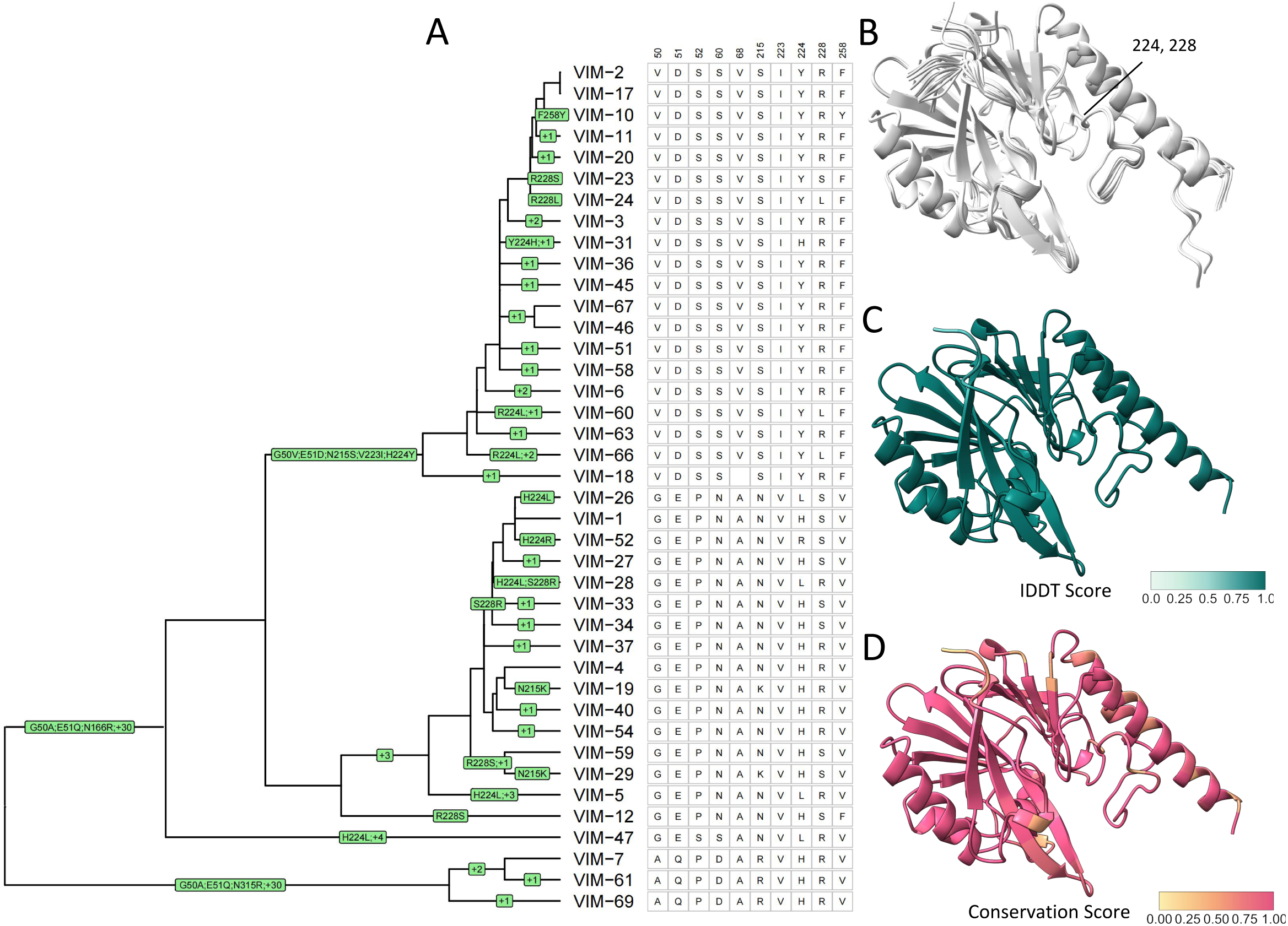
Key amino acid substitutions in VIM variants map to structurally conserved enzymes. (A) Guide tree inferred from multiple sequence alignment with annotated amino acid variation. Key residues associated with antimicrobial resistance profiles are highlighted at positions 50, 51, 52, 60, 68, 215, 223, 224, 228, and 258. (B) Structurally aligned AlphaFold2-predicted protein models for all VIM variants. (C) Residue-level lDDT scores calculated using FoldMason structural alignment, reflecting structural conservation. (D) Amino acid conservation scores derived from residue similarity and the bio3d substitution matrix.

Phylogenetic analysis showed that VIM variants formed distinct evolutionary branches. VIM-1/VIM-4-like and VIM-2-like variants constituted the main framework. VIM-1- and VIM-4-related variants were closely related, whereas VIM-2 and VIM-7 formed more independent lineages, indicating clear subclade differentiation (Figure 6A).

Despite multiple amino acid substitutions among variants, the overall protein structure was highly conserved. AlphaFold2-based structural prediction showed that all VIM variants maintained a canonical MBL fold with high confidence (mean pLDDT = 96.0, SD = 7.903). The catalytic core also showed nearly identical spatial conformation across variants (Figures 6B and 6C). These findings suggest that the VIM family is constrained by strong structural and functional requirements during long-term evolution.

Sequence-conservation analysis further showed that amino acid substitutions were not randomly distributed. Instead, they were concentrated at a limited set of positions, including residues 50, 51, 52, 60, 68, 215, 223, 224, 228, and 258 (Figure 6D). Conservation scores at these sites were lower than the full-sequence average, suggesting that they represent hypervariable regions in VIM evolution. Residues 224 and 228 are located near the active site and underwent repeated substitutions across multiple phylogenetic branches. These patterns imply potential roles in fine-tuning substrate recognition or enzymatic activity while maintaining catalytic stability.

Collectively, VIM variants showed an evolutionary pattern characterised by a highly conserved core structure and continuous diversification at localised residues. Under the constraint of a stable metal-binding and catalytic framework, sequence diversity appeared to arise mainly through repeated substitutions at a limited number of hotspots. These patterns identify residues that may represent candidate sites for adaptive diversification and warrant further functional investigation (Figure 6).

### 7. Healthcare-associated environmental reservoirs contribute to *bla*_VIM_ circulation

To explore the ecological distribution of *bla*_VIM_-positive strains, we analysed isolate sources across clinical, environmental, and animal compartments. (Figure 7A and Supplementary Data 1). Human-derived isolates were dominant (2,873, 95.7%), followed by environmental isolates (98, 3.3%) and animal isolates (approximately 21, 0.7%). These data indicate that *bla*_VIM_ currently circulates mainly in clinical contexts. Among environmental isolates, sample types were unevenly distributed (Figure 7B). Wastewater-related samples were the most common, followed by hospital environmental samples such as surfaces and equipment. Other environmental sources were less frequent. This pattern suggests that environmental *bla*_VIM_ is concentrated in niches closely linked to healthcare activity.

**Figure 7.**
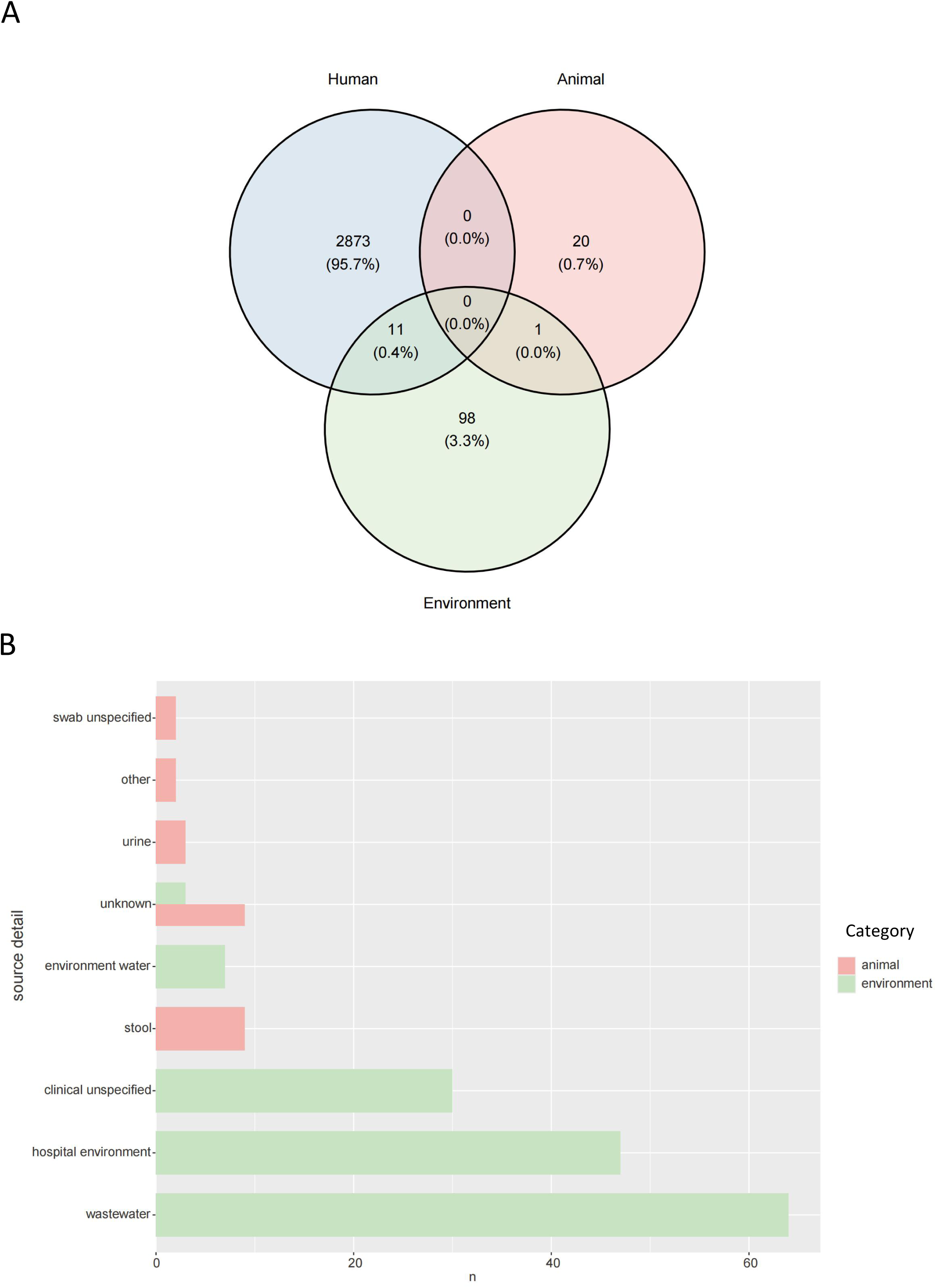
Host origins of *bla*_VIM_-positive genomes across human, animal, and environmental niches. (A) Venn diagram showing the overlap of defined VIM clusters among human, animal, and environmental genomes, reflecting possible cross-niche clonal transmission, shared genetic contexts, or independent isolation. (B) Counts of genomes recovered from specific non-human sources. Source data are provided in the Source Data file.

Animal isolates were limited in number but diverse in origin, mainly comprising faecal and related samples (Figure 7B). All major *bla*_VIM_ subtypes were detected in animals, with *bla*_VIM-2_ relatively more common. Host sources included avian, mammalian, and aquatic species, although counts were small within each group.

Cross-source analysis identified several VIM clusters shared across ecological compartments (Figure 7A). Eleven VIM clusters were detected in both environmental and human sources, and a small number overlapped between environmental and animal sources. These findings indicate limited but detectable sharing of *bla*_VIM_-positive lineages or genetic contexts between clinical and non-clinical compartments, suggesting potential connectivity between healthcare-associated reservoirs and broader ecological niches.

## Discussion

Using 5,617 *bla*_VIM_-positive genomes collected worldwide between 1999 and 2025, this study this study provides a global genomic view of the host, clonal, plasmid, and mobile-element contexts underlying VIM dissemination. The principal finding is that global *bla*_VIM_ spread does not appear to follow a single universal transmission pathway. Instead, two contrasting host-associated patterns were evident: in *P. aeruginosa*, particularly for *bla*_VIM-2_, dissemination was strongly associated with successful high-risk clones, whereas in *Enterobacterales*, *bla*_VIM-1_ and *bla*_VIM-4_ were distributed across diverse species through broad-host-range plasmids. Moreover, widespread geographic dissemination was concentrated in a limited number of successful VIM–host–vector combinations, while many others remained restricted to individual countries or regions. The results indicate that *bla*_VIM_ has evolved from a resistance determinant associated with localised European healthcare outbreaks into a globally disseminated and highly interconnected resistance network^7, 69^. Together, these findings support a host-dependent and hierarchical transmission architecture in which clonal expansion, plasmid-mediated horizontal transfer, and mobile genetic elements contribute at different levels to the persistence and geographic spread of *bla*_VIM_.

*bla*_VIM-1_, *bla*_VIM-2_, and *bla*_VIM-4_ accounted for the great majority of VIM-positive isolates, indicating that the global burden of *bla*_VIM_ is driven by a limited number of successful variants rather than uniform expansion of the entire VIM family. This distribution is broadly consistent with previous regional and global observations^7, 8^. However, our temporal analysis further showed a shift in the relative predominance of the two major variants: *bla*_VIM-1_ was slightly more frequent before 2019, whereas *bla*_VIM-2_ subsequently became the most frequently detected variant. This shift suggests a transition from a traditional VIM-1-dominated pattern associated with *Klebsiella* and other *Enterobacterales* toward a VIM-2-dominated pattern centred on *P. aeruginosa*. Thus, changes in global VIM subtype composition may reflect not only the success of individual resistance alleles but also the expansion of the bacterial populations and genetic platforms with which they are associated. This succession may reflect changing antimicrobial selection pressure. The widespread use of ceftazidime-avibactam and related agents targeting KPC-type carbapenemases may indirectly favour intrinsically resistant MBL-producing strains. At the same time, VIM-2-associated high-risk *P. aeruginosa* clones have strong environmental adaptability and nosocomial transmission capacity, which may further support their global expansion^70–72^.

Compared with the more clone-dominated transmission often described for *bla*_KPC_ or *bla*_OXA-48_, *bla*_VIM_ displays a multilayered transmission architecture^70^. *bla*_VIM-1_ and *bla*_VIM-4_ spread mainly among *Enterobacterales* through broad-host-range plasmids such as IncA, IncC, IncHI2A, and IncL/M. By contrast, *bla*_VIM-2_ is closely coupled to high-risk *P. aeruginosa* clones, including ST111, ST235, and ST233. This dual mode, combining plasmid-driven horizontal transfer with stabilisation in successful clones, probably contributes to the long-term persistence of *bla*_VIM_^24, 73–76^. Among *Enterobacterales*, broad-host-range plasmids enable repeated cross-species transfer and expansion of the resistance network. Some *bla*_VIM_-associated plasmids also carry multiple resistance determinants and complex recombination structures. These features suggest that modular recombination and hybrid plasmid formation may enhance host range and persistence^77–79^. In *P. aeruginosa*, by contrast, *bla*_VIM_ dissemination appears to rely more on sustained clonal expansion and stable chromosomal or lineage-specific contexts than on classical broad-host-range plasmid transfer^80–82^. Thus, the relative contribution of clonal and plasmid-mediated transmission differs markedly between major bacterial hosts.

The transmission pattern of *bla*_VIM-2_ in *P. aeruginosa* was particularly distinct from that observed in *Enterobacterales*. Many *bla*_VIM-2_-positive genomes lacked canonical replicons or showed chromosomal localisation, implying greater dependence on clonal expansion and long-term persistence than on frequent plasmid transfer. ST235 and ST111 are established global high-risk multidrug-resistant *P. aeruginosa* clones carrying multiple resistance and virulence determinants^71, 72^. Our analysis further suggests that these clones act as core reservoirs for *bla*_VIM-2_ and also carry diverse *bla*_VIM_ subtypes, with ST235 showing exceptional subtype diversity. High-risk *P. aeruginosa* clones may therefore function not only as amplifiers but also as stable genetic backgrounds for long-term *bla*_VIM_ maintenance and diversification. Stable chromosomal integration of resistance genes can promote long-term maintenance, while environmental persistence of *Pseudomonas*-associated resistance elements may further support prolonged circulation in hospital settings^82, 83^.

A further important finding was that plasmid-mediated dissemination did not necessarily correspond to uniform global spread. Although several broad-host-range plasmids crossed bacterial species and national boundaries, most VIM variant–plasmid cluster combinations remained confined to individual countries or regions. Only a limited subset showed cross-country or intercontinental dissemination. This pattern suggests that global *bla*_VIM_ spread is driven disproportionately by a small number of successful plasmid–variant combinations, whereas many other transmission events remain geographically restricted. Such a pattern is consistent with a model in which regional transmission predominates, with only selected genetic platforms achieving wider international dissemination.

Mobile genetic elements also played a central role in *bla*_VIM_ dissemination. More than 96% of *bla*_VIM_-positive regions were associated with integrative genetic elements, many of which co-occurred with insertion sequences or phage-related elements to form complex nested structures. Such multilayered genetic architectures may enhance *bla*_VIM_ mobility, recombination, and long-term maintenance across different bacterial backgrounds^84^. Previous studies have established integrons as key platforms for the long-term dissemination and accumulation of MBL genes^3, 85^. ISCR elements, ICE-like regions, and transposons can further facilitate resistance-gene capture and mobilisation^86–92^. Our data indicate that different *bla*_VIM_ subtypes may be associated with distinct mobile-element contexts. *bla*_VIM-1_ was preferentially associated with insertion sequence/integrative genetic element contexts, whereas *bla*_VIM-2_ was more frequently linked to phage/integrative genetic element contexts. This subtype-specific modularity suggests that different VIM variants may persist within different mobile-element architectures during long-term dissemination^19, 93–96^.

VIM also shows an important association with healthcare-related environmental reservoirs. Environmental isolates in this study were concentrated in hospital wastewater, sinks, and drains, and several VIM clusters were shared between human and environmental sources^30, 33, 97, 98^. These findings support hospital water systems as persistent reservoirs for VIM-positive bacteria. Previous studies have shown that VIM-positive *P. aeruginosa* can colonise sinks and drainage systems for prolonged periods through biofilm formation. Therefore, in addition to patient-to-patient transmission, VIM may also be maintained through a patient-hospital environment-patient cycle^30, 32, 99^. Persistent environmental colonisation increases outbreak risk and creates opportunities for horizontal gene transfer among bacterial communities, particularly in hospital wastewater^100, 101^.

Although *bla*_VIM_-positive isolates were identified across human, environmental, and animal sources, human-derived isolates accounted for the great majority of the present dataset. Animal isolates were rare, whereas environmental isolates were mainly associated with healthcare settings. Therefore, our results provide stronger support for circulation between clinical and healthcare-associated environmental compartments than for extensive transmission across the broader One Health interface. Increasing evidence suggests that carbapenem resistance genes have moved beyond clinical systems into broader ecological networks, including healthcare environments, wastewater, natural water bodies, and animal hosts^32, 102^. Discharge of hospital wastewater into natural environments may facilitate further dissemination of *bla*_VIM_. Continuous gene exchange between environmental and clinical bacteria may also accelerate long-term ecological circulation. These findings indicate that *bla*_VIM_ control should extend beyond clinical infection management. Surveillance of hospital wastewater, aquatic environments, and other non-clinical reservoirs should be incorporated into One Health strategies^103, 104^.

Our structural analysis provides additional insight into the evolution of the VIM family. Although amino acid substitutions were observed across variants, overall protein structures remained highly conserved, consistent with previous structural studies^10, 11^. Repeated substitutions at residues near the active site, particularly positions 224 and 228, nevertheless suggest that adaptive evolution occurs through localised functional tuning rather than global structural remodelling^12, 14^. Because these residues have previously been shown to influence substrate recognition and carbapenem hydrolysis efficiency, the clustering of hypervariable sites around functionally relevant regions may reflect ongoing adaptation of VIM enzymes^10^. With the increasing clinical use of cefiderocol and aztreonam-avibactam, continued monitoring is needed to determine whether emerging VIM variants acquire altered substrate profiles that compromise current salvage therapies^37, 105, 106^. However, the present study did not experimentally assess enzyme activity or substrate specificity, and the functional consequences of these substitutions therefore require further biochemical validation.

This study has several limitations. First, reliance on public genomic data introduces geographic and temporal sampling bias. Countries with high sequencing capacity may be over-represented, whereas low- and middle-income settings may be under-represented. In addition, temporal changes in isolate numbers may partly reflect changes in sequencing and surveillance intensity rather than true changes in population prevalence. Second, most genomes were derived from short-read assemblies, limiting the resolution of complex mobile-element structures and complete plasmid architecture. Third, the small number of environmental and animal isolates may underestimate the contribution of non-clinical reservoirs to global *bla*_VIM_ transmission and limits broader One Health inference. Fourth, the present analysis did not reconstruct the precise temporal origin or directionality of major transmission events, which will require dedicated time-scaled phylogenetic and phylogeographic analyses. Finally, incomplete genomic and metadata coverage for rare VIM variants limited our ability to fully characterise the transmission patterns of low-frequency subtypes, and the functional significance of candidate sequence substitutions remains to be experimentally validated.

In summary, *bla*_VIM_ has evolved from a regionally recognised resistance determinant into a complex, globally disseminated resistance network. Its spread is shaped by interactions among high-risk clones, broad-host-range plasmids, and multilayered mobile genetic elements. In *P. aeruginosa*, *bla*_VIM_ dissemination is characterised mainly by clonal stabilisation and persistence within successful high-risk lineages, whereas in *Enterobacterales* it is driven more strongly by plasmid-mediated cross-species transfer. Importantly, most VIM–host–vector combinations appear to remain regionally restricted, while only a limited number achieve wider geographic dissemination. These findings support a host-dependent and hierarchical transmission model for *bla*_VIM_ and highlight the need for targeted genomic surveillance of successful bacterial clones, transferable plasmid platforms, and healthcare-associated environmental reservoirs.

## Acknowledgements

This study was financially supported by the Scientific Research Foundation of the State Key Laboratory of Vaccines for Infectious Diseases and Xiang An Biomedicine Laboratory (Grant No. 2025XAKJ0100004), the Zhongnanshan Medical Foundation of Guangdong Province (ZNSXS 20260088), the Tertiary Education Scientific research project of Guangzhou Municipal Education Bureau (Grant No. 2024312107), and the Guangdong Provincial Medical Research Foundation (Grant No. A2025209). All funding agencies were not involved in the study design, experimental implementation, data acquisition and analysis, result interpretation, manuscript drafting and revision, as well as the submission decision of the present work.

---

**Figure S1.**
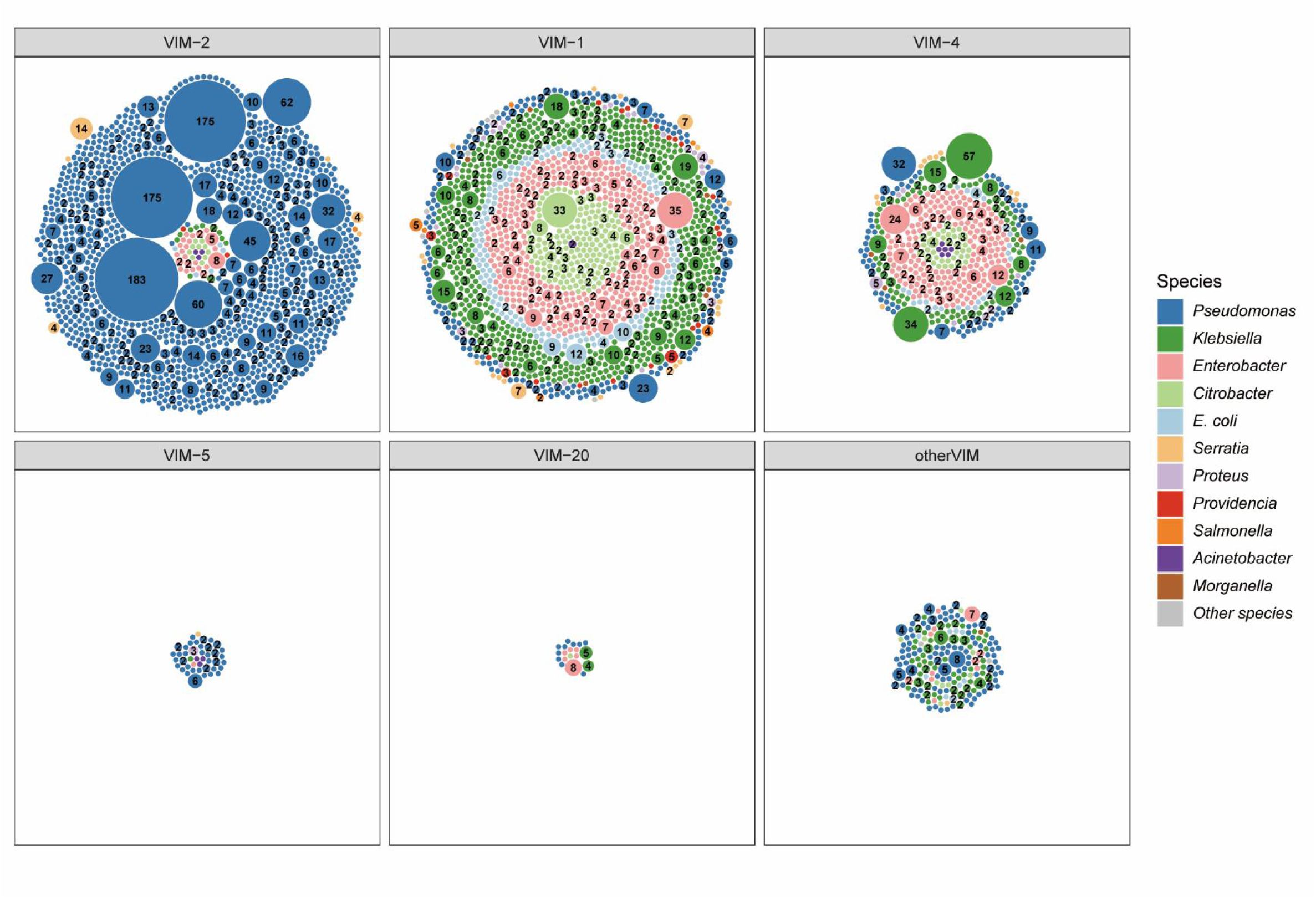
Size distribution of VIM clusters in the dataset. Bubble size and internal numerical labels represent the genome count of each VIM cluster. Clusters containing only one genome are unlabeled.

**Figure S2.**
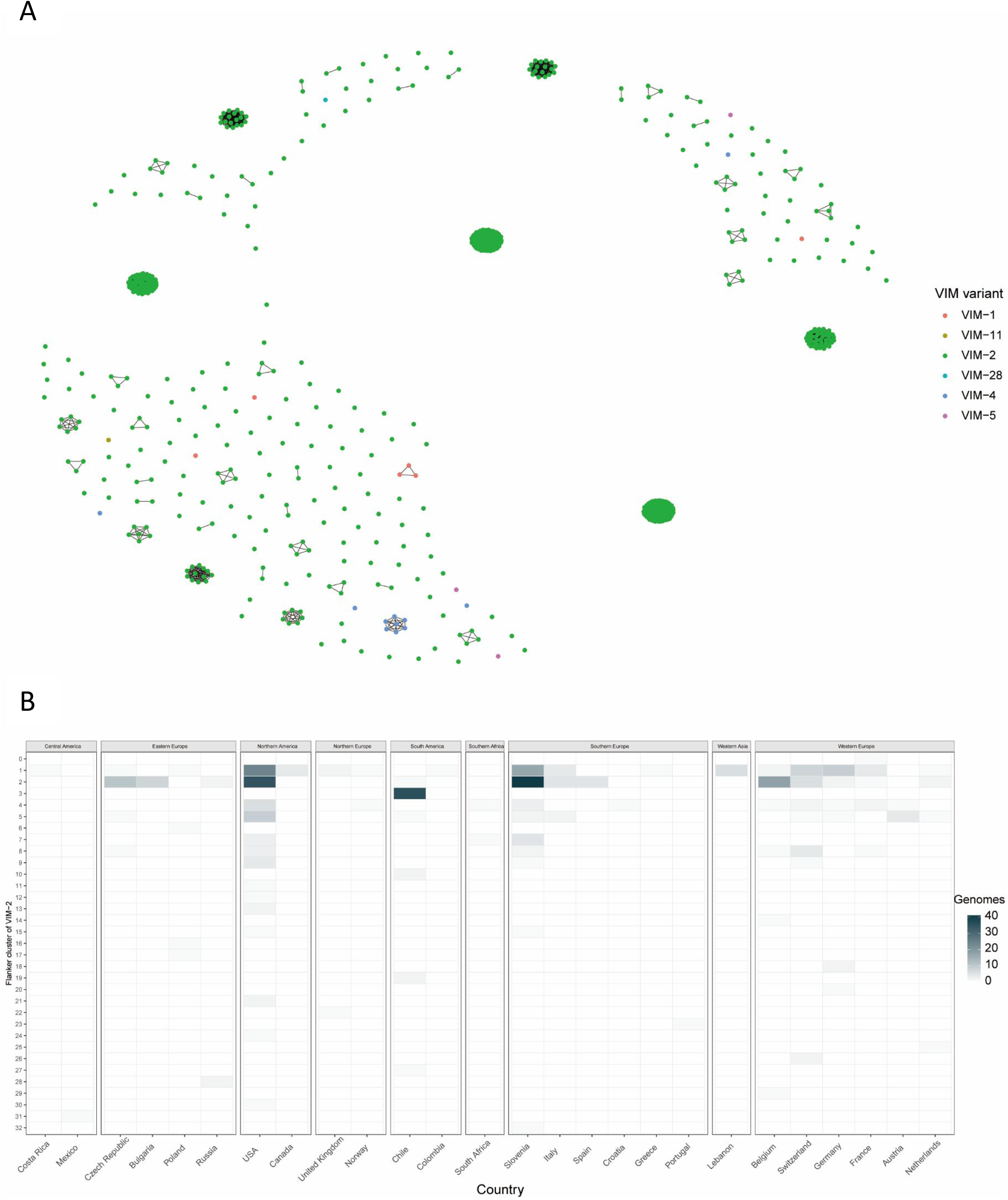
Global dissemination and genetic diversity of *bla*_VIM_-carrying *P. aeruginosa* ST111. (A) Network of clonally related ST111 genomes colored according to *bla*_VIM_ variant. (B) Heatmap showing region-specific independent acquisition of the target β-lactamase gene among *bla*_VIM_-positive ST111 strains.

**Figure S3.**
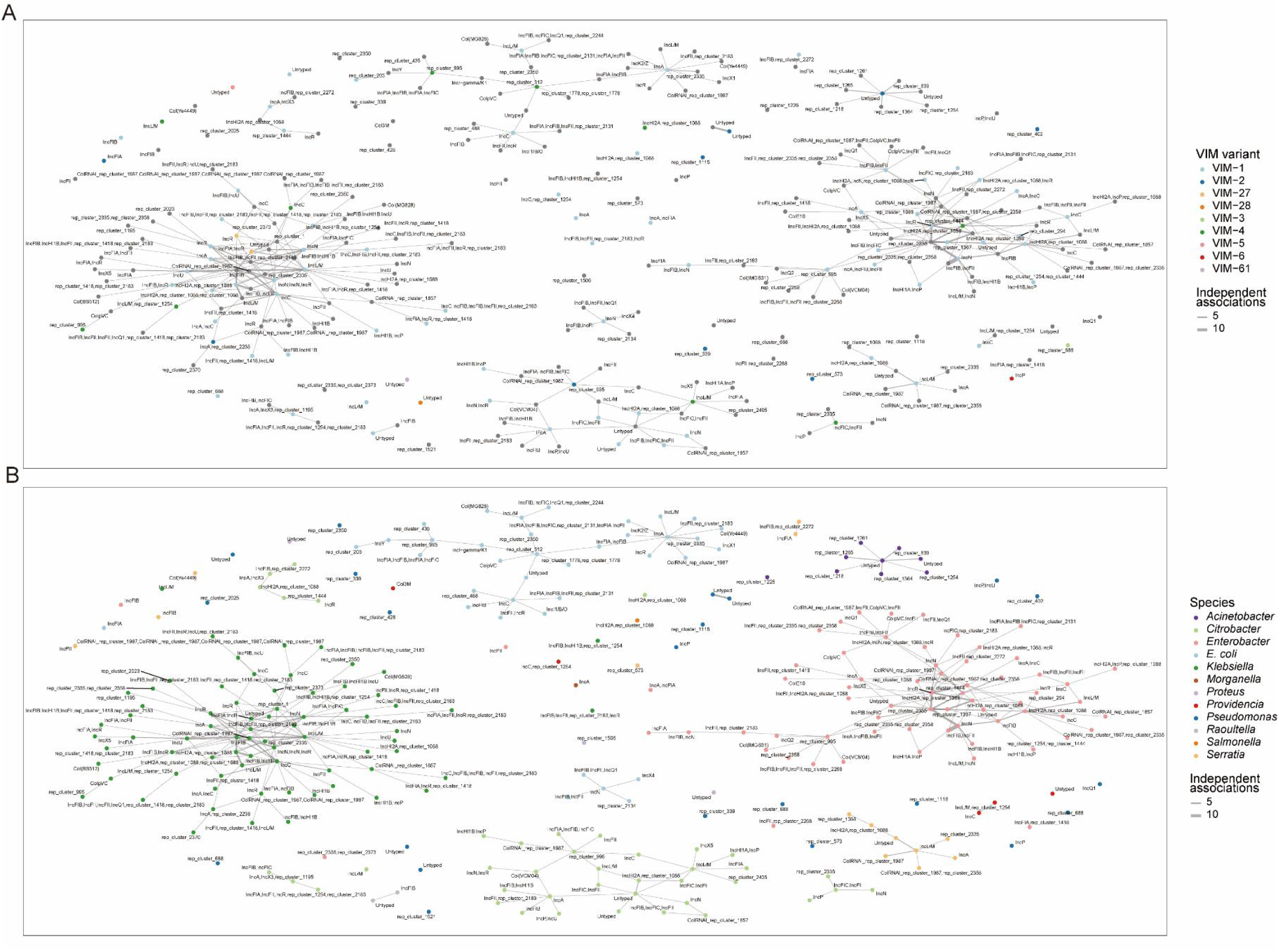
Co-occurrence network of *bla*_VIM_-harboring plasmids and non-*bla*_VIM_ plasmids. (A) Network coloured by *bla*_VIM_ variant. (B) Network coloured by bacterial species. Genomes were dereplicated into VIM clusters to reduce clonal redundancy before analysis. Edges connect plasmids detected in the same genome, with edge thickness proportional to co-occurrence frequency.

**Figure S4.**
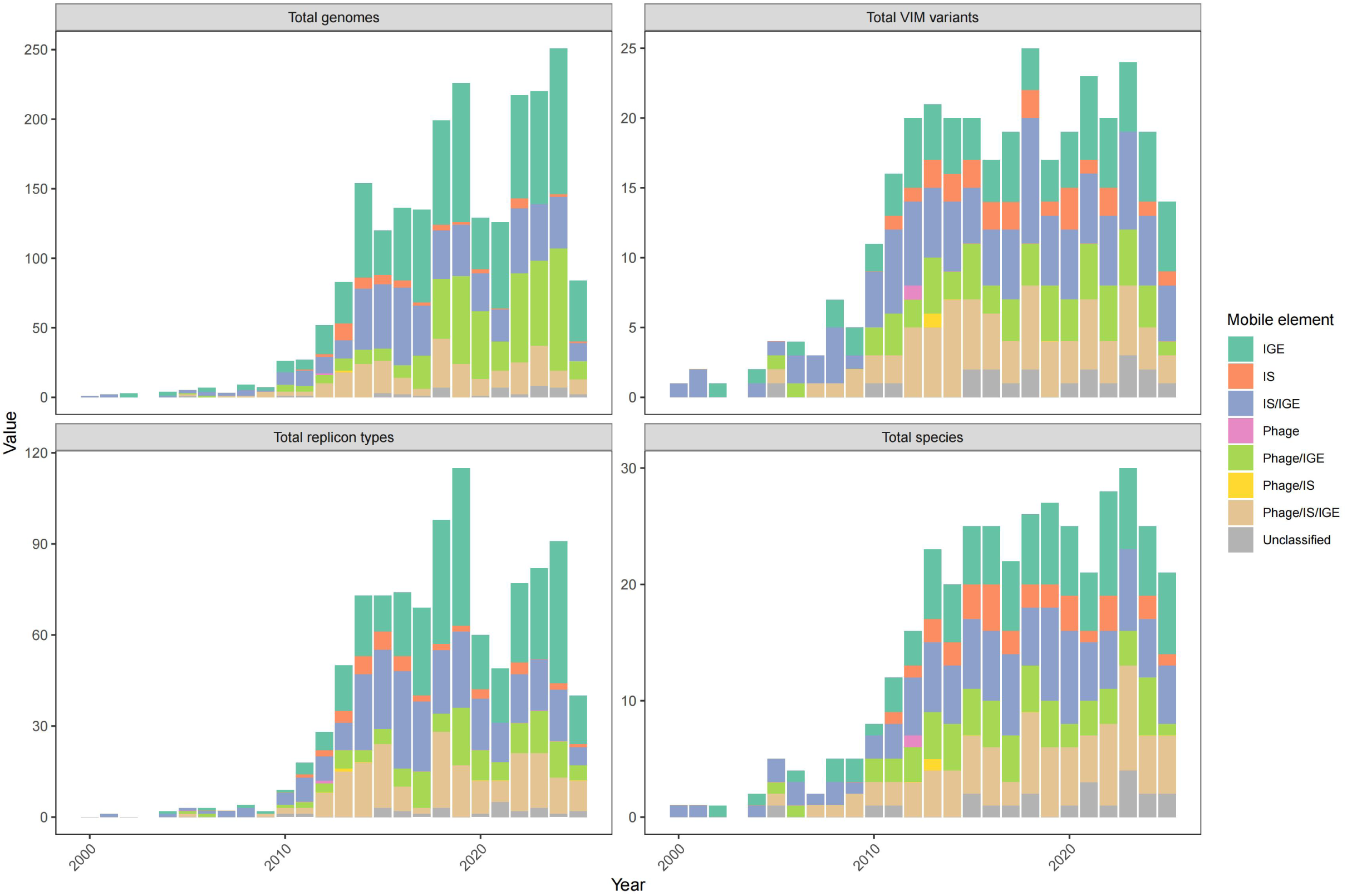
Temporal association of mobile genetic elements with VIM carbapenemases. Cumulative bar chart showing the composition of linked mobile elements and their distribution across bacterial species, *bla*_VIM_ variants, total genomes, and plasmid types. Raw data are provided in Supplementary Data 1.

Description of additional supplementary files

File Name: Supplementary Data 1

Description: metadata table used in this study.

File Name: Supplementary Data 2

Description: supporting information for VIM-1, VIM-2, and VIM-4 analyses in Figure 2.

File Name: Supplementary Data 3

Description: pLDDT scores predicted by AlphaFold2 and ColabFold.

